# The genetic architecture of lipoprotein(a): simplicity, complexity, and mutability

**DOI:** 10.64898/2026.11.16.767202

**Authors:** Ronen E. Mukamel, Simone Rubinacci, Nolan Kamitaki, Robert E. Handsaker, Margaux L.A. Hujoel, David Tang, Steven A. McCarroll, Po-Ru Loh

## Abstract

High levels of lipoprotein(a) (Lp(a)) increase risk of cardiovascular diseases. Lp(a) concentration is primarily determined (*>*80%) by genetic variation at *LPA*, particularly by its large, size-polymorphic kringle IV-2 (KIV-2) repeat. However, because variation of the KIV-2 repeat is difficult to measure, the precise ways that genetic variants at *LPA* and other loci determine Lp(a) levels have been unclear, especially in individuals of non-European ancestry. Here, we developed new computational techniques to accurately genotype KIV-2 variation in whole-genome-sequenced population cohorts. Applying these methods to UK Biobank (n=490,414) and All of Us (n=245,394) provided insights into the genetic determinants of Lp(a) and the mutational dynamics of the KIV-2 repeat, which we found expands or contracts intra-allelically once every *∼*500 generations. Variation in Lp(a) levels among people of all genetic ancestries was accurately predicted by a single, transferable genetic model capturing *cis*-epistatic effects of KIV-2 length and protein-altering polymorphisms in *LPA*. Controlling for predicted effects of *LPA* alleles revealed an additional, highly polygenic contribution to Lp(a) variation: *LPA*-adjusted Lp(a) levels associated with genetic variation at over 300 loci (*P<*5*×*10^−8^), much like other blood lipids. These results elucidate the genetic architecture and mutational origins of Lp(a) variation, showing how detailed characterization of a largely monogenic trait can provide surprising insights for complex trait genetics.

## Main

Lipoprotein(a), like low-density lipoprotein (LDL), is a cholesterol-carrying blood lipid, high levels of which increase risk of cardiovascular diseases including myocardial infarction, ischemic stroke, and aortic stenosis^1–5^. Lp(a) levels vary dramatically across individuals, from *<*0.1 to *>*200 mg/dL (*∼*500 nmol/L)^6^. This variation, which is *∼*80–90% genetically determined by genetic variation at the *LPA* locus^7–9^, has fascinated lipidologists and geneticists for decades. The primary determinant of Lp(a) concentration, responsible for roughly half of its variance in human populations, is the length of the kringle IV-2 (KIV-2) repeat in the *LPA* gene^7, 10–12^. *LPA* encodes apolipoprotein(a) (apo(a)), which binds an LDL-like particle to form Lp(a), and its KIV-2 repeat (consisting of *∼*1–40 copies of a 5.5-kb, 2-exon repeat unit) encodes an allelically variable number of kringle domains within apo(a)^13, 14^. This size variation affects the efficiency of apo(a) secretion from hepatocytes: larger apo(a) isoforms encoded by longer KIV-2 repeats take longer to transit the endoplasmic reticulum, resulting in lower plasma Lp(a) levels^15, 16^. Beyond KIV-2 size, several single-nucleotide polymorphisms in *LPA* (including variants within the KIV-2 repeat) cause loss of function or mis-splicing of *LPA*, reducing Lp(a) levels in the European populations in which these variants are common^17–26^.

While the main genetic drivers of Lp(a) variation among European-ancestry individuals are now recognized, several key questions about the genetics of Lp(a) are unanswered. Even at the *LPA* locus, the genetic architecture of Lp(a) in non-European ancestries is largely uncharacterized^27^. Lp(a) genetics in European versus non-European ancestries is known to be sufficiently different for SNP-based genetic predictors to be non-transferable^28, 29^, but whether this can be explained solely by population differences in allele frequencies and linkage disequilibrium patterns of causal variants has been unclear^11^. Beyond *LPA*, knowledge of polygenic contributors to Lp(a) variation is sparse: unlike other blood lipids, for which hundreds of genetic risk loci have been identified by genome-wide association studies (GWAS)^30^, only 39 GWAS loci have been identified for Lp(a)^24^. Additionally, little is known about the mutational dynamics of the KIV-2 repeat: how it expanded from a single repeat unit within an ancient gene duplication of the *PLG* gene^13^, and how it continues to expand and contract today^7, 31^.

The main challenge that has impeded further characterization of Lp(a) genetics is the difficulty of accurately measuring haplotype-resolved KIV-2 repeat variation—by far the largest effect on Lp(a) concentration—in large, diverse population cohorts^27^. We previously developed a population-genetic approach to this problem, using haplotype sharing within large cohorts to resolve sequencing coverage-based estimates of total (“diploid”) KIV-2 repeat copy number into estimates of allele-specific copy number; however, this approach requires many closely or distantly related individuals to achieve high accuracy^26^. Recently, a novel approach based on haplotype-informative sequence variants within the KIV-2 repeat was shown to accurately estimate allele-specific copy numbers from whole genome sequencing (WGS) data for heterozygous individuals^32^. Here, we combined these two ideas to further improve KIV-2 genotyping accuracy in the UK Biobank (UKB)^33^ and All of Us (AoU)^34^ cohorts, enabling new insights into KIV-2 copy-number mutation and the genetic architecture of Lp(a).

### Deep haplogroup structure among rarely-recombining KIV-2 repeat alleles

To inform our approach to estimating KIV-2 repeat lengths and to learn about the evolutionary history of human KIV-2 repeat alleles, we first examined an ancestral recombination graph previously inferred for haplotypes from the 1000 Genomes Project Phase 3 (1KGP3) and archaic hominins^35^. The coalescent tree at the KIV-2 region exhibited a deep split approximately 1.8 million years ago (Fig. 1A). Each branch of this split (segregating with rs6902102:G*>*A) was represented at high frequency across 1KGP3 populations (derived AF=0.27–0.51; Fig. 1A and Supplementary Fig. 1).

**Figure 1.**
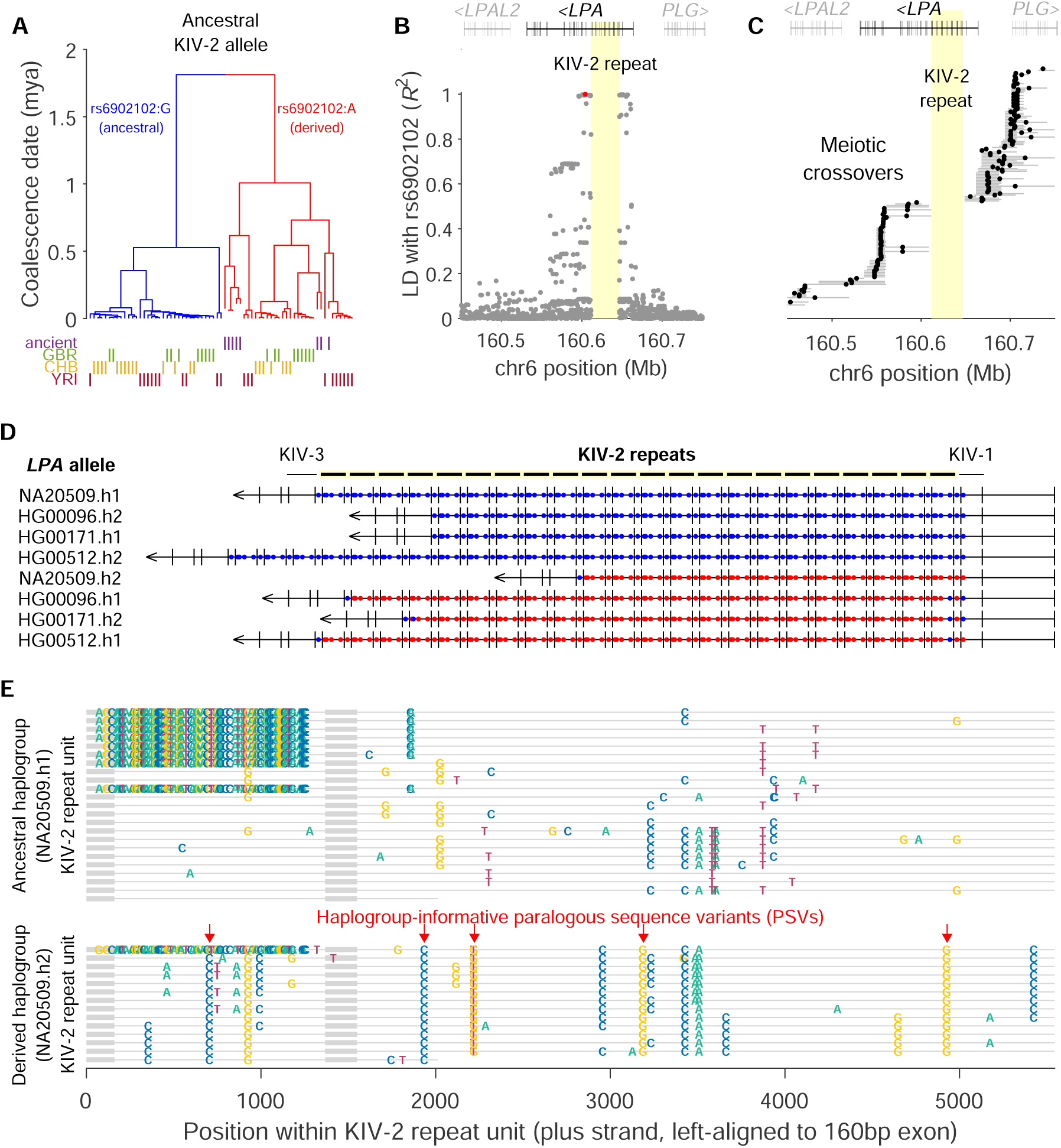
*LPA* KIV-2 repeat alleles segregate into haplogroups that rarely recombine. **(A)** Coalescent tree of modern human and archaic hominin haplotypes inferred by Relate^35^. The branches of the deepest split (blue/red) segregate with a SNP haplotype containing rs6902102. **(B)** Linkage disequilibrium in 1KGP3 (n=2,504) between rs6902102 (red) and other SNPs at the *LPA* locus. Positions, GRCh38 coordinates. **(C)** Meiotic crossovers at the *LPA* locus resolved to *<*100 kb in the deCODE data set^38^. Dots, averages of start and end positions. **(D)** Haplogroup-informative paralogous sequence variants within assembled KIV-2 repeat alleles^39^. Each KIV-2 repeat unit contains five dots (one per PSV) colored blue or red according to the allele of the PSV within that repeat unit. Alleles from the rs6902102:A haplogroup (bottom four alleles containing primarily red dots) exhibit incomplete intra-allelic homogeneity (blue dots in the first and last repeat units). **(E)** Visualization of all single-nucleotide PSVs observed within two KIV-2 repeat alleles. The PSV-dense region in the short intron distinguishes type A and type B repeats^6^.

Linkage disequilibrium (LD) analysis in 1KGP3 (n=2,504) showed that rs6902102 tags a long haplotype spanning the KIV-2 repeat that has rarely been broken by recombination (Fig. 1B). The unusually low recombination rate within the KIV-2 repeat—one recombination per *∼*300,000 meioses based on LD in 1KGP3^36, 37^, hundreds of times lower than the genome-wide average for a *∼*100-kb region—was corroborated by data from meiotic crossovers in parent-offspring pairs (126,427 meioses)^38^: among the 73% of crossovers that had been resolved to *<*100 kb resolution, none overlapped the KIV-2 repeat (Fig. 1C and Supplementary Note 1).

Several SNPs flanking the KIV-2 repeat co-segregated with rs6902102 (Fig. 1B)—and thus the two deeply-diverged KIV-2 repeat haplogroups (Fig. 1A)—suggesting the possibility that other such SNPs might have accrued within the KIV-2 repeat itself, potentially early enough to homogenize across repeat alleles. Analyses of short-read WGS data in UKB and assembled KIV-2 repeat alleles^39^ identified five such paralogous sequence variants (PSVs) that exhibited near-perfect intraallelic homogeneity across the repeat units of each KIV-2 allele (Fig. 1D,E, Supplementary Fig. 2, and Supplementary Table 1), including the two haplogroup-informative PSVs recently described^32^. For four of the five PSVs, alleles in the derived haplogroup (rs6902102:A) commonly exhibited incomplete intra-allelic homogeneity, but usually only because the first or the last repeat unit of the allele carried the reference PSV allele, whereas all other repeat units carried the alternate PSV allele (Fig. 1D, Supplementary Fig. 2, and Supplementary Table 1). These four PSVs presumably originated as mutations along the rs6902102:A branch of the initial split between KIV-2 alleles (Fig. 1A) and either occurred on a very short, ancestral KIV-2 allele containing a single KIV-2 repeat unit or were homogenized across the repeat by subsequent gene conversions, expansions, and contractions that did not involve meiotic recombination. Only 2 potentially recombinant KIV-2 alleles (with haplogroup-discordant flanking SNPs) were observed among 5,008 alleles in 1KGP3 (Supplementary Note 1).

### Estimating allele-specific KIV-2 repeat lengths using haplogroup-informative PSVs

We leveraged the five haplogroup-informative PSVs to accurately estimate KIV-2 repeat lengths from short-read WGS data, building on ideas from the DRAGEN KIV-2 caller^32^ and our previous population-genetic approach^26^. The DRAGEN KIV-2 caller uses two haplogroup-informative PSVs to partition diploid KIV-2 copy number (estimated from WGS read-depth) into allele-specific repeat lengths based on PSV allele fractions^32^. This innovative approach has both strengths and limitations compared to the population-genetic approach^26^: while the PSV-based approach is only applicable to heterozygous individuals with one allele from each haplogroup (comprising approximately half of the human population), it does not require information from other individuals who co-inherited a matching KIV-2 allele. As such, it can provide accurate length estimates of KIV-2 alleles carried by only one or a few individuals in a cohort (e.g., alleles carrying recent copy-number mutations or present in minority ancestries).

Incorporating population-genetic ideas into PSV-based KIV-2 copy-number phasing considerably improved allele-sizing accuracy while retaining the strengths of the DRAGEN approach. In brief, we used a haplotype-sharing model to impute, for each PSV, the number of repeat units within a KIV-2 repeat allele that deviated from the expected PSV allele. These deviations were easily imputed from SNP haplotypes because they tended to occur in the relatively stable first and last repeat units of KIV-2 alleles (Fig. 1D and Supplementary Figs. 2 and 3). Being able to account for these deviations enabled all five haplogroup-informative PSVs to be used for KIV-2 copy-number partitioning. This new technique, together with a new method for precisely calibrating WGS read-depth (Methods and Supplementary Notes 2 and 3), allowed us to generate highly accurate PSV-based KIV-2 allele length estimates for 215,936 UKB participants (RMSE=0.4 repeat units based on analyses of sibling pairs in UKB with both haplotypes identical by descent (IBD2); Fig. 2A,B); in comparison, the DRAGEN KIV-2 caller had RMSE=0.7 repeats in benchmarks using 1000 Genomes Project (1000G) samples^32, 40^ (Supplementary Fig. 4 and Supplementary Note 4). The PSV-based estimates were broadly concordant with estimates of KIV-2 lengths that we generated for the full UKB cohort using our previous haplotype-sharing-based approach^26^.

**Figure 2.**
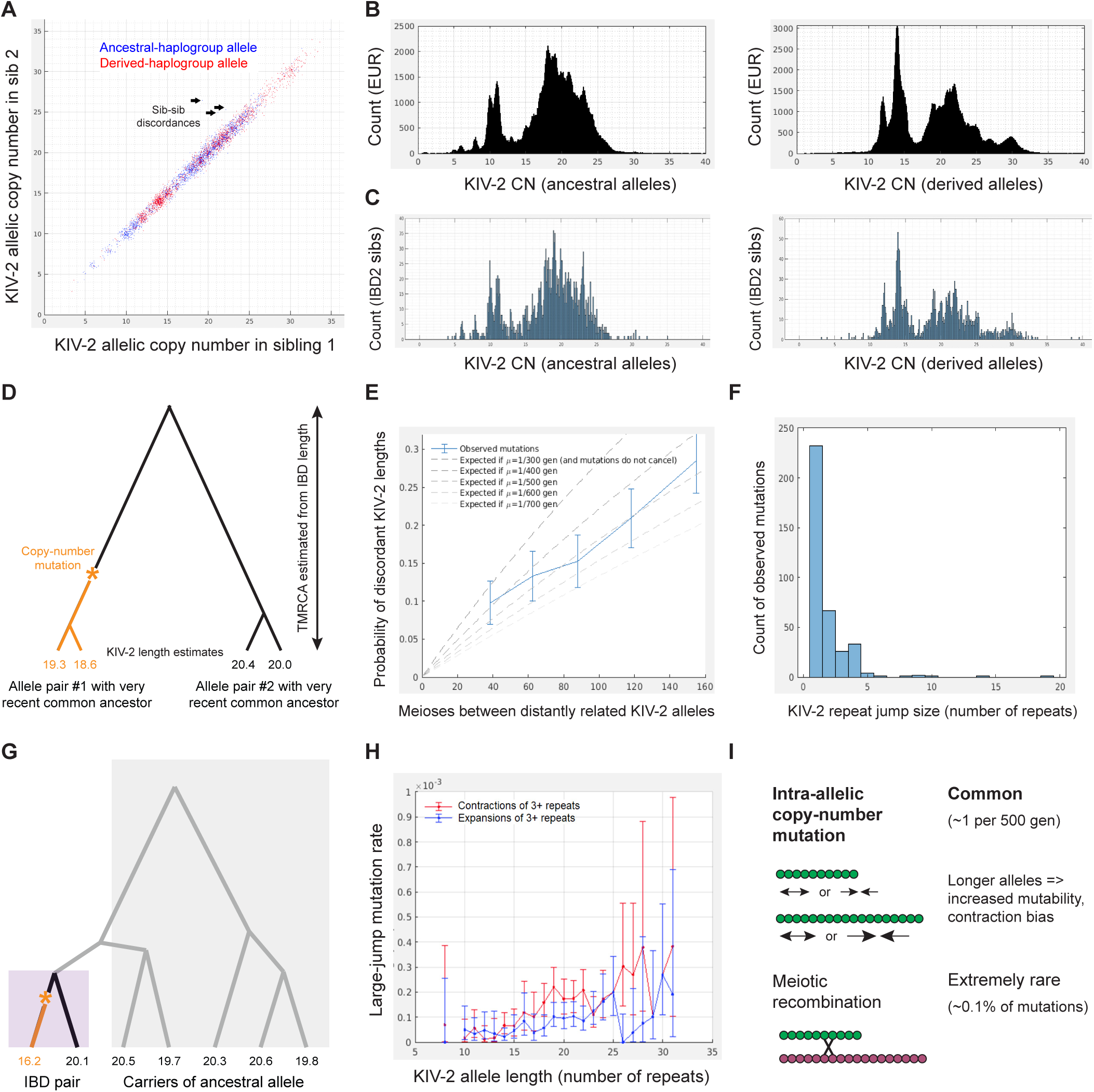
Mutational dynamics of KIV-2 repeat expansion and contraction inferred from KIV-2 mutations in IBD tracts. **(A)** Comparison of allele-specific KIV-2 repeat length estimates in pairs of siblings in the UKB cohort. Data are shown for sibling pairs ascertained to be IBD2 at *LPA* and heterozygous for the two KIV-2 haplogroups (i.e., carrying one allele each from the ancestral rs6902102:G and derived rs6902102:A haplogroups). Allele-specific KIV-2 length estimates were obtained using the PSV-based approach to phase WGS-derived diploid copy number estimates. Callouts indicate allele length discordances between siblings, presumably reflecting de novo mutations. Five alleles with lengths exceeding the plot boundaries (38–47 repeat units) are not shown. **(B)** Histograms of PSV-based KIV-2 allele length estimates in EUR-ancestry UKB participants heterozygous for the two haplogroups. Left, ancestral alleles; right, derived alleles. **(C)** Histograms of refined allele length estimates obtained by averaging two independent PSV-based measurements of the same allele inherited by two IBD2 siblings. Combining two measurements of an allele’s length from two related individuals reduces measurement error to the extent that the distribution of copy-number estimates exhibits discrete integer modes, enabling many alleles to be resolved to exact copy numbers with high confidence. **(D)** Schematic of approach used to estimate KIV-2 copy-number mutation rate. Among alleles that could be confidently sized by combining independent estimates from two related individuals, we identified pairs of allleles sharing more-distant IBD (time to most recent common ancestor (TMRCA) up to *∼*100 generations, estimated from IBD length) and modeled the probability of observing discordant allele lengths as a function of TMRCA and mutation rate (*µ*). **(E)** Observed and predicted fractions of IBD pairs with discordant allele lengths as a function of the number of meioses *m* separating the alleles (*∼*2*×*TMRCA). The data are consistent with a mutation rate *µ* of 1 mutation per *∼*500 generations. Importantly, the fraction of discordant pairs increases roughly linearly with TMRCA, indicating a minimal effect of spurious discordances arising from measurement errors (which would inflate discordance rates for all IBD pairs equally, independent of TMRCA). **(F)** Histogram of copy-number differences observed between pairs of confidently-sized KIV-2 alleles belonging to to IBD pairs. This distribution reflects the jump-size distribution of KIV-2 expansions and contractions (up to a small amount of noise introduced by measurement error and by rare instances of IBD lineages containing multiple mutations). Jump sizes are absolute values (i.e., expansions are not distinguished from contractions) because the source allele was unknown in this analysis. **(G)** Schematic of approach used to estimate allele-specific rates of large expansions and contractions. We analyzed rates of large discordances (*≥*3 repeat units) between lengths of alleles with a recent common ancestor, determining the source allele using imputation^41^. **(H)** Estimated rates of large expansions and contractions (*≥*3 repeat units) for KIV-2 alleles of different lengths. Data are plotted for allele lengths for which IBD lineages descended from alleles of that length span a total of *>*10,000 meioses. **(I)** Graphical summary of KIV-2 mutational dynamics. Error bars, 95% CIs in all panels.

### Mutational dynamics of KIV-2 repeat expansion and contraction

Having accurate length estimates for hundreds of thousands KIV-2 alleles allowed us to investigate several longstanding questions about the mutational dynamics of the KIV-2 repeat^6^: how frequently it expands and contracts, whether expansions and contractions tend to involve few or many repeat units, and how allele length affects mutability. Only one example of a KIV-2 copy-number mutation has previously been documented: an intra-allelic loss of 9 repeats, observed once among 376 meioses^7, 31^. Discordances between KIV-2 allele lengths in IBD2 sibling pairs in UKB corroborated the occasional occurrence of multi-unit mutations: among *∼*2,500 IBD2 sibling pairs (representing *∼*10,000 meioses) with PSV-based KIV-2 length estimates, three large discordances (of approximately 3, 4, and 7 repeat units) were observed (Fig. 2A). However, this analytical approach could not assess the frequency of 1- or 2-unit mutations (which were within the tail of measurement error) or distinguish between expansions and contractions.

We devised two population-genetic strategies to overcome these limitations (Supplementary Note 5). First, to enable detecting copy-number mutations of all sizes (including 1-unit changes), we identified a subset of *∼*10,000 alleles for which we could confidently call exact KIV-2 lengths by combining concordant estimates of the allele’s length from two related individuals (Fig. 2C). We then examined rates of discordances among pairs of these confidently-sized alleles that were descended from a more-distant ancestor up to *∼*100 generations ago (Fig. 2D). Analyzing this data set (representing a total of *∼*200,000 meioses) showed that copy-number mutations within KIV-2 alleles occur roughly once per 500 generations (Fig. 2E)—three orders of magnitude more frequently than meiotic recombination—and usually involve small changes of 1–2 repeat units (Fig. 2F).

Second, we performed a separate IBD-based analysis in which we distinguished between expansions and contractions by imputing source alleles^41^ (Fig. 2G). To obtain the sample size needed here, we restricted our attention to allele length discordances of 3 or more repeat units (which could be confidently ascertained from individual PSV-based allele length measurements). This analysis indicated that longer KIV-2 alleles are more prone to contraction than expansion and more mutable overall (Fig. 2H).

Taken together, these results characterize the dynamics of frequent intra-allelic expansions and contractions of the KIV-2 repeat (Fig. 2I), providing clues for further understanding the mutational mechanism (probably unequal sister chromatid exchange, as at centromeres^42^) that generates abundant copy-number variation at highly mutable macrosatellite repeats^43–45^.

### Predicting lipoprotein(a) levels from variation at *LPA* across genetic ancestries

Despite the large proportion of variation in Lp(a) levels attributable to KIV-2 length variation and other genetic variation at the *LPA* locus^7–9^, modeling and predicting Lp(a) levels from genotypes has been surprisingly difficult. SNP-based predictors have achieved accuracy (*R*^2^) up to 0.6 in populations of European (EUR) genetic ancestry^22, 28, 46–49^ but have exhibited low transferability to other ancestries^28, 29^ (*R*^2^*<*0.03 in UKB participants of African (AFR) or East Asian (EAS) ancestry; Fig. 3A). To understand this phenomenon and build an accurate, portable genetic model of Lp(a), we analyzed phased *LPA* haplotypes (including WGS-derived KIV-2 allele lengths and SNPs and indels in KIV-2) together with Lp(a) measurements for 411,680 unrelated UKB participants.

**Figure 3.**
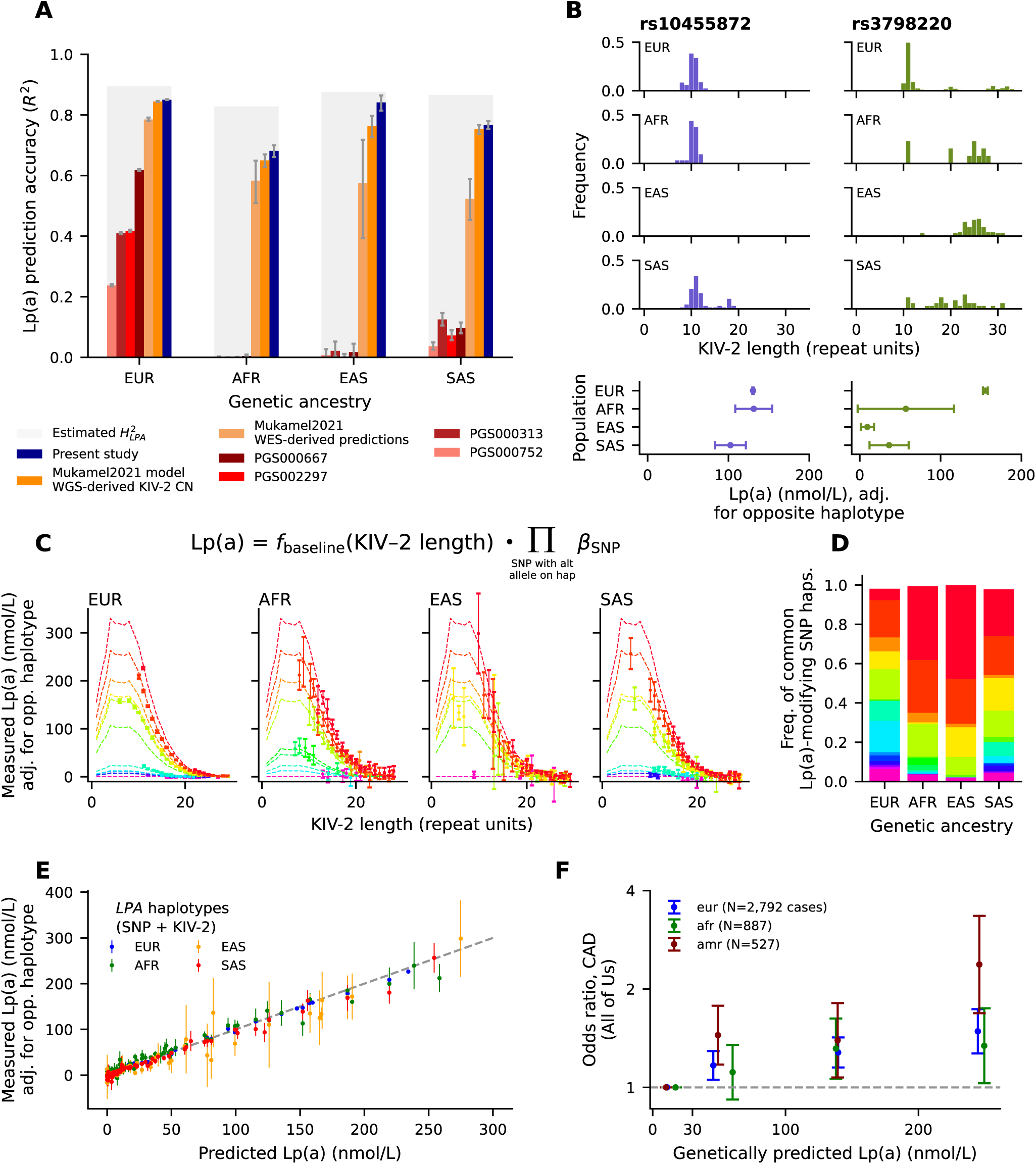
Predicting Lp(a) from genetic variation at *LPA* across ancestries. **(A)** Lp(a) prediction accuracy (*R*^2^ between predicted and measured Lp(a)) per genetic ancestry in UKB for: SNP-based polygenic scores (shades of red, listed by PGS Catalog ID); our previous prediction model^26^ (shades of orange); and the model presented in this study (blue). Accuracy of our previous model is shown using either WES-derived *LPA* SNP+KIV-2 haplotypes^26^ (light orange) or haplotypes generated from WGS data in this study (dark orange). Heritability attributable to the *LPA* locus (grey shading) was estimated from Lp(a) measurements in siblings (Supplementary Note 7). All predictors were benchmarked in unrelated individuals with PSV-phased KIV-2 lengths; predictions using WES-derived haplotypes (light orange) were further restricted to individuals in the initial UKB WES release. **(B)** (top) KIV-2 allele length distributions (in PSV-phased individuals) for rs10455872*>*G and rs3798220*>*C haplotypes, stratified by genetic ancestry. Whereas *LPA* alleles carrying rs10455872*>*G have short KIV-2 repeats in all ancestries, this is true of rs3798220*>*C only in EUR. (bottom) Mean Lp(a) contributed by rs10455872*>*G and rs3798220*>*C haplotypes per genetic ancestry. Lp(a) measurements were adjusted for the predicted contribution of the opposite haplotype. **(C)** Predicted Lp(a) (dashed line) and measured Lp(a) (dots) per KIV-2 allele length, stratified by common Lp(a)-modifying SNP haplotype (colors; each color corresponds to a SNP combination with AF*>*1% in at least one genetic ancestry; null alleles are grouped together in the bottom category). Measured Lp(a) was adjusted for the opposite haplotype’s predicted contribution. Each plot (corresponding to an ancestry) displays data for SNP+KIV-2 haplotypes with frequency *>*0.1% in that ancestry. KIV-2 allele lengths were estimated using PSV-based phasing and rounded to the nearest integer. **(D)** Frequencies of the common Lp(a)-modifying SNP haplotypes displayed in panel **C** per ancestry. **(E)** Measured vs. predicted Lp(a) contributed by the SNP+KIV-2 haplotypes displayed in panel **C**, stratified by genetic ancestry. **(F)** Odds ratio for coronary artery disease (CAD) per tranche of median predicted Lp(a) in AoU, stratified by genetic ancestry. Participants were binned according to their predicted Lp(a) (bin cutoffs indicated by *x*-tick marks). Odds ratios (from logistic regression) are relative to individuals with predicted Lp(a)*<*30 nmol/L. Error bars, 95% CIs.

Examining the haplotypes of two common SNPs (rs10455872 and rs3798220) that were among the first to be associated with high Lp(a) levels^2^ shed light on the challenges: the extent to which these SNPs tagged short KIV-2 alleles varied across populations^50^ (Fig. 3B and Supplementary Note 6), consistent with the combined effects of recombination and intra-allelic KIV-2 mutation. Moreover, *cis*-epistatic interactions among variants on a haplotype that modify its Lp(a) production^26^ exacerbated the difficulty of SNP-based prediction (Supplementary Note 6).

To overcome these challenges, we modeled the amount of Lp(a) produced by each of an individual’s two *LPA* haplotypes as the product of a baseline contribution, determined by the length of the KIV-2 repeat, with a multiplicative effect of each Lp(a)-modifying coding or 5’ UTR variant present on the haplotype (Fig. 3C). This simple approach achieved high accuracy across EUR, AFR, EAS, and South Asian (SAS) genetic ancestries (*R*^2^=0.85, 0.68, 0.84, and 0.77, respectively), explaining 96%, 82%, 96%, and 89% of Lp(a) heritability attributable to the *LPA* locus in these ancestry groups (Fig. 3A and Supplementary Note 7). The high prediction accuracies reflected two improvements compared to our previous analysis based on whole-exome sequencing (WES) data from 49,959 UKB participants^26^. First, simply replacing WES-derived KIV-2 and SNP/indel haplotypes with more accurate, WGS-derived haplotypes (within our previous model using KIV-2 length and 23 fine-mapped coding and 5’ UTR SNPs in *LPA*^26^) substantially increased prediction accuracy (Fig. 3A). Second, we augmented our previous model with four additional, ancestry-specific coding variants in *LPA* (EUR allele frequency (AF)*≤*0.01%; AF*≥*0.3% in another ancestry) that were identified by conditional analyses (Supplementary Fig. 5) and associated with complete loss of Lp(a) production. Incorporating these variants into an improved Lp(a) model (Fig. 3C and Supplementary Tables 2 and 3) further increased prediction accuracy (Fig. 3A), particularly in the EAS group, in which a common variant (EAS AF=1.1%) disrupts a canonical splice donor site of KIV-2 exon 1^51^ on short KIV-2 alleles that would otherwise produce high Lp(a).

The high accuracy of our Lp(a) model suggests that the genetic architecture of Lp(a) is shaped primarily by a small set of causal variants in *LPA* with consistent effects across genetic ancestries. The amount of Lp(a) attributable to each common *LPA* haplotype (defined by KIV-2 allele length and SNPs used in the model) was accurately predicted by the model across genetic ancestries (Fig. 3C,E). The major differences in Lp(a) genetics across ancestries^11, 27–29, 50, 52^ are thus attributable almost entirely to differences in the frequencies of causal variants with *cis*-epistatic effects (Fig. 3D, Supplementary Tables 2 and 3, and Supplementary Note 8) rather than by ancestry-dependent effect sizes.

To further validate the transferability of this Lp(a) predictor and explore its potential utility for cardiovascular risk stratification in diverse cohorts, we applied the model to WGS and coronary artery disease (CAD) data for 230,573 AoU participants. Higher predicted Lp(a) associated increased CAD risk across all well-represented ancestries (increased CAD log-odds of 17% (95% CI 11–24%), 13% (4–22%), and 39% (24–56%) per 100 nmol/L increase in predicted Lp(a) in EUR, AFR, and Admixed American (AMR) ancestries, respectively; Fig. 3F and Supplementary Note 9).

### Polygenic influences of genetic variation at 331 loci on Lp(a) levels

Although Lp(a) levels are mainly determined by *LPA* haplotypes, the *APOE*-*ɛ*2 allele reduces Lp(a) levels^22, 53, 54^, and GWAS in UKB have identified up to 37 other loci associated with Lp(a)^24, 29, 55^, hinting at complex genetic modulation beyond *LPA*. We reasoned that controlling for the predicted Lp(a) production of an individual’s two *LPA* alleles might greatly increase power to observe such effects. In UKB, *LPA*-adjusted Lp(a) was heritable (*h*_g_^2^=0.25 (0.01)) and associated with genetic variants at 331 loci (*P<*5*×*10^−8^), similar to other blood lipids^30^ (Fig. 4A, Supplementary Table 4, and Supplementary Note 10). Index variants at most of these loci (87%) also associated with at least one other lipid level (*P<*0.01 for LDL, HDL, total cholesterol, or triglycerides; Fig. 4B–D and Supplementary Table 4).

**Figure 4.**
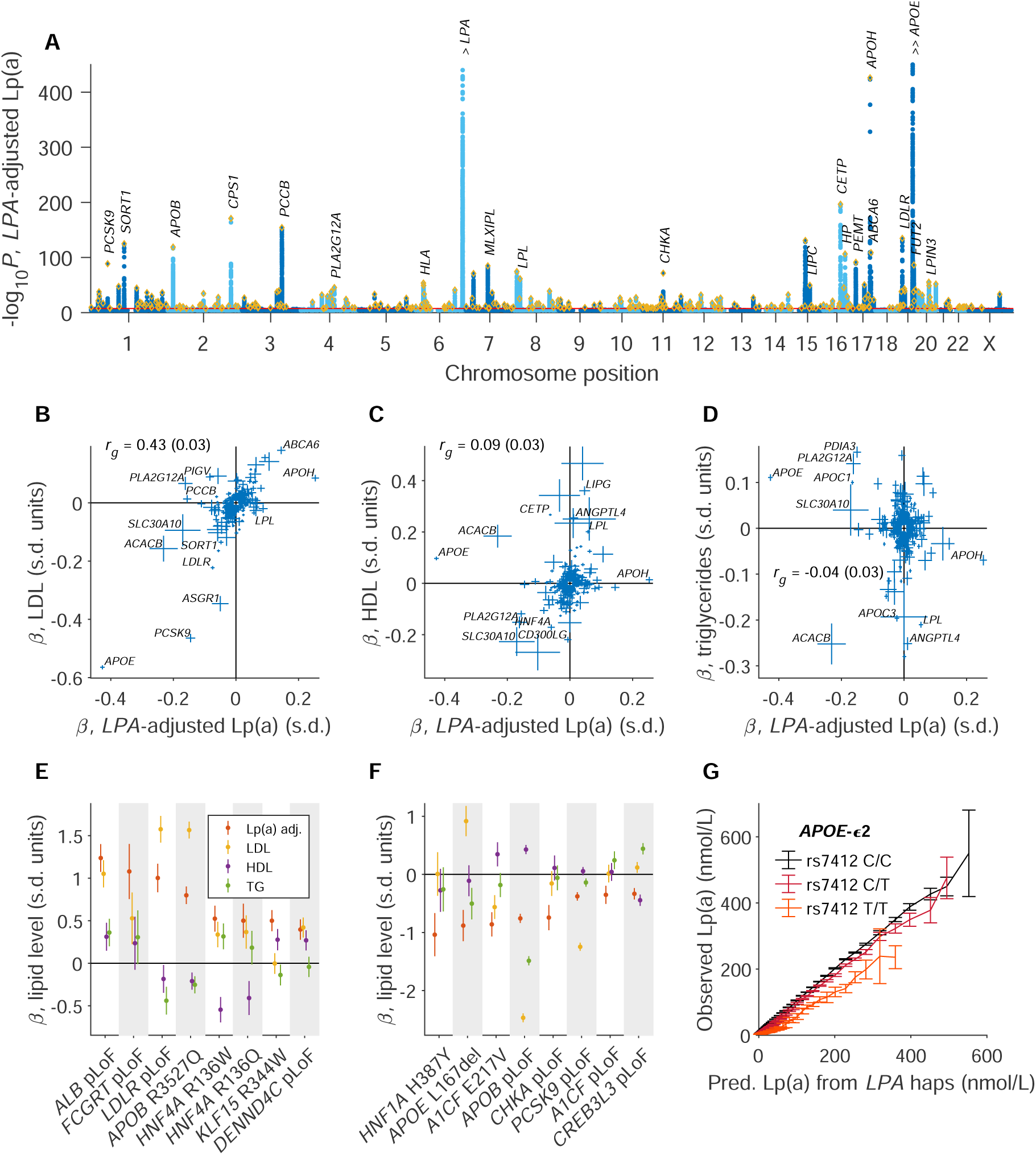
Genome-wide genetic influences on lipoprotein(a) levels. **(A)** Associations of common and low-frequency variants (MAF*>*0.1%) with Lp(a) levels adjusted for predictions based on *LPA* diplotypes. Gold diamonds, lead variants. Among the top 30 loci, loci with putative target genes are labeled. Association strengths at two loci exceed the upper limit of the plot: *APOE* (*P*=6.3*×*10^−3061^) and *LPA* (*P*=5.6*×*10^−943^, reflecting remaining genetic effects at *LPA* not captured by our model). **(B,C,D)** Scatter plots comparing effect sizes (*β*) of common and low-frequency variants on *LPA*-adjusted Lp(a) (*x*-axis) and other lipid levels (*y*-axis; LDL, HDL, and triglycerides in panels **B-D**). Units, standard deviations. Genetic correlations (*r*_g_) are indicated within the plots. **(E,F)** Effect sizes (*β*) of rare coding variants on *LPA*-adjusted Lp(a) and other lipid levels. Lp(a)-increasing effects are in panel **E**; Lp(a)-decreasing effects, panel **F**. **(G)** Complex effects of the *APOE*-*ɛ*2 allele (rs7412) on Lp(a). Error bars, 95% CIs in all panels.

*LPA*-adjusted Lp(a) had high genetic correlation with LDL (*r*_g_=0.43 (s.e. 0.03); Fig. 4B), consistent with Lp(a) being composed of an LDL particle bound to apo(a)^6^. However, some genetic variants associated with opposing effects on LDL and *LPA*-adjusted Lp(a) (Fig. 4B), including the *PLA2G12A* D111N missense variant, the common *LPL* stop gain haplotype rs328, and variants associated with expression of *PCCB* (which encodes a subunit of an enzyme involved in lipid catabolism) and *PIGV* (which encodes an enzyme involved in glycolipid synthesis). At 92 loci, the index variant’s normalized effect size for *LPA*-adjusted Lp(a) was at least twice that of any other lipid level (Supplementary Table 4). Genes at these loci were enriched in the synthesis of phosphatidylcholine pathway (*P*=1.43*×*10^−5^, Bonferroni-adjusted *P*=0.034). Two of the 28 genes in this pathway were previously associated with Lp(a) levels (*PEMT* and *CHKA*^24^); seven more were implicated here (*LPIN3*, *PHOSPHO1*, *MFSD2A*, *ACHE*, *STARD10*, *CHKB*, and *CHPT1*).

Protein-coding variants appeared to drive many genetic associations with *LPA*-adjusted Lp(a), like other lipids^56^, including 9 of the top 15 common-variant associations reported here (Supplementary Table 4). Rare coding variants (AF*<*0.1%) in 13 genes associated with strong effects on Lp(a) (Fig. 4E,F and Supplementary Tables 5 and 6). As one example, variants predicted to cause loss of function (pLoF) of *ALB*, which encodes albumin, a serum protein involved in cholesterol transport, associated with a 1.24 (0.08) s.d. increase in *LPA*-adjusted Lp(a) (Fig. 4E), corresponding to a median increase in Lp(a) of 66% relative to predictions based on *LPA* diplotypes (Supplementary Fig. 6). Most rare coding variants affected multiple lipid levels, though with varying effect sizes and directions; in particular, the *APOE* L167del inframe deletion associated with strongly increased LDL^57^ but 47% decreased Lp(a) (*P*=8.1*×*10^−13^; Fig. 4F and Supplementary Fig. 6). Two effects of rare coding variants were specific to Lp(a): Lp(a)-lowering effects of *CHKA* pLoF variants and a missense variant in *HNF1A*, which encodes a transcription factor active in hepatocytes with a binding site in the *LPA* promoter^58^ (Fig. 4F). These variants associated with 37% and 48% reductions in Lp(a), respectively (*P*=7.1*×*10^−12^ and *P*=8.3*×*10^−9^; Supplementary Fig. 6). Common and rare variant effects appeared to act roughly multiplicatively on Lp(a) levels predicted from *LPA* diplotypes (Supplementary Figs. 6 and 7). *APOE*-*ɛ*2 genotype exhibited a complex, non-additive effect on Lp(a), as also observed for other lipids^59^ (Fig. 4G and Supplementary Fig. 7). These results show that beyond the large effect of *LPA* haplotypes on Lp(a) production, Lp(a) levels are also shaped by genome-wide influences in a manner similar to other lipid measurements, suggesting common mechanisms affecting Lp(a) regulation or catabolism.

### Hundreds of rare protein-altering variants in *LPA* generate null or hypomorphic alleles

Accurately modeling effects of common variation at *LPA* on Lp(a) levels additionally enabled a well-powered screen for rare variants in *LPA* that associated with large deviations from predicted Lp(a), suggesting strong effects of these variants on apo(a) function. Previous data sets have offered limited power to undertake such analysis: our WES-based analysis of 10% of the UKB cohort identified six such variants (five in *LPA* exons or splice regions and one in the 5’ UTR) with EUR AF 0.01%–0.09%^26^, including one previously reported^23^. Here, analyzing WGS data for the full UKB cohort using new statistical methods enabled evaluating variants that were as much as 20-fold rarer (AF=0.0005%; 5 carriers in UKB). We considered 742 such rare variants (AF between 0.0005% and 0.1%) within exons, splice regions, and the promoter of *LPA*—including SNPs and indels within the KIV-2 region (which comprised 49% of these variants)—which we genotyped and phased in 394,896 unrelated EUR-ancestry UKB participants with available Lp(a) measurements. To assess each variant’s association with Lp(a) and estimate its multiplicative effect size on an allele’s Lp(a) production, we developed a likelihood ratio test that accounted for the heteroskedasticity of Lp(a) measurements and the amounts of Lp(a) predicted to be produced by each individual’s two alleles (Supplementary Note 11).

This analysis identified 322 rare variants that associated with Lp(a) at an FDR*<*0.05 threshold, among which 157 variants (i) reached a stringent *P<*10^−5^ significance threshold, (ii) associated with large effects on an allele’s Lp(a) production (*±*50% or more), and (iii) associated independently of other variants (Fig. 5A–C and Supplementary Table 7). These 157 variants exhibited strong enrichment in functional categories predicted from DNA sequence (Fig. 5A,B), similar to variant effect distributions seen in exome-wide association studies^60, 61^. The variants included 42 pLoF variants, 11 putative cryptic splice variants (SpliceAI^62^ delta score*≥*0.2), and 95 missense variants. All but one variant associated with strongly reduced Lp(a) (Fig. 5C and Supplementary Fig. 8), and the lone association with increased Lp(a) appeared likely to be a false positive (Supplementary Fig. 8). Although these variants affected only a small fraction of all *LPA* haplotypes (combined EUR AF=0.01), they associated with large individual-level reductions in Lp(a). The large majority of the 157 variants appeared likely to causally alter apo(a): only 7 of the 157 likely-causal variants were among the 291 tested variants with no predicted protein-altering impact. Even among these 7 non-coding variants, 2 variants associated with complete loss of an allele’s Lp(a) production and appeared to regulate *LPA* transcription or translation (Fig. 5C): one is a promoter variant that disrupts a conserved HNF-1*α* binding motif (chr6:160664317*>*A, *P*=2.9*×*10^−47^; Supplementary Fig. 9) and the other is a previously identified 5’ UTR variant^26^.

**Figure 5.**
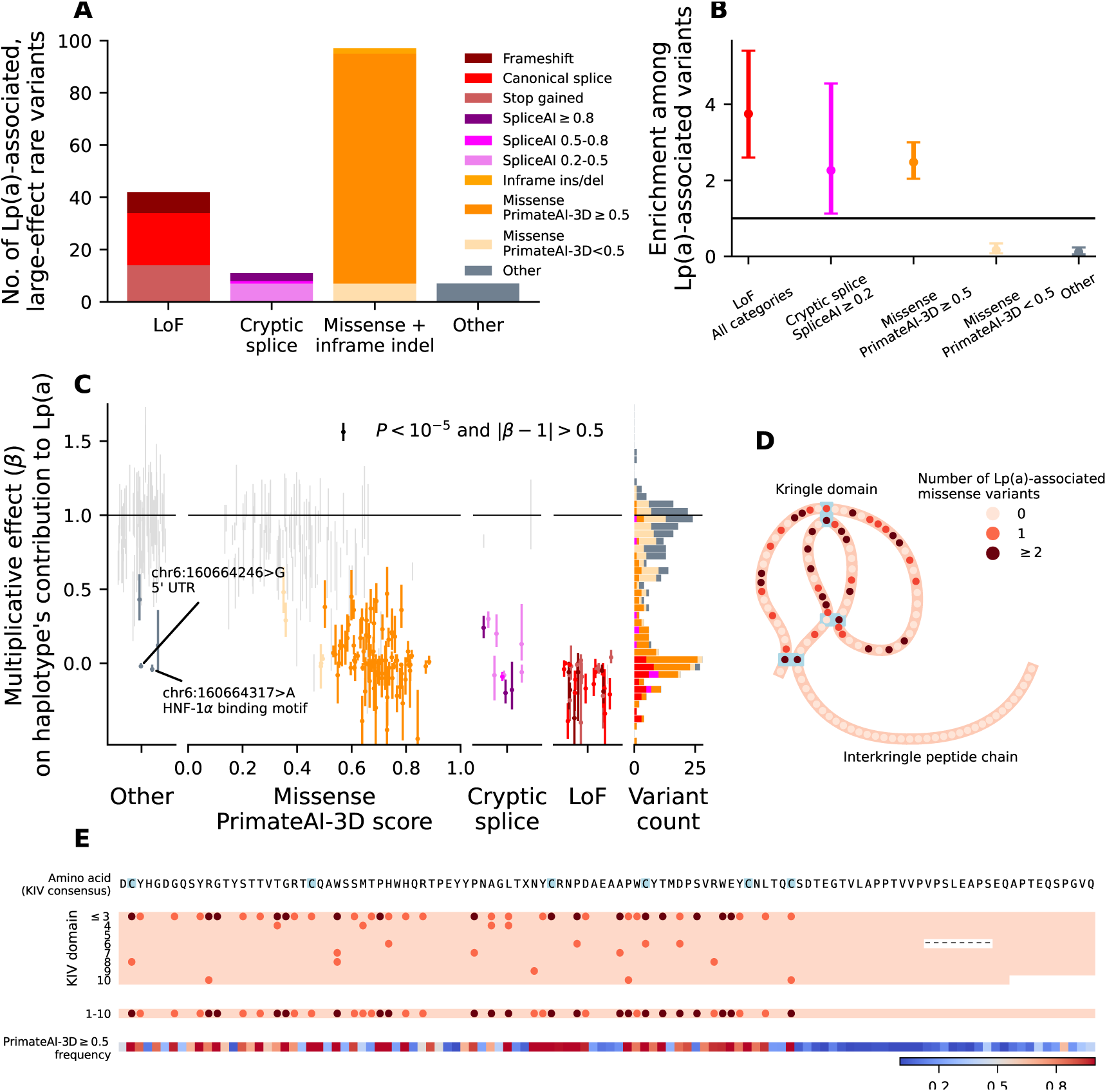
Hundreds of rare protein-altering variants in *LPA* generate null or hypomorphic alleles. **(A)** Distribution of predicted functional consequences of rare variants in *LPA* (AF=0.0005%–0.1%) that associated (*P<*10^−5^) with a large effect on an allele’s Lp(a) production (*±*50% or more) and that passed LD-filtering. **(B)** Enrichment (per functional category) of Lp(a)-associated variants tabulated in panel **A** relative to tested variants. Error bars, 95% CIs. **(C)** (left) Maximum likelihood estimate of each rare variant’s multiplicative effect on a haplotype’s contribution to Lp(a). (right) Stacked histogram of multiplicative effect sizes, grouped by functional annotation (colored as in panel **A**). On the left side, variants not determined to have likely-causal large effects on Lp(a) are plotted in light grey. Error bars, approximate 95% CIs. Variants with CI span *>*0.5 are omitted for clarity; full data are plotted in Supplementary Fig. 8. **(D)** Schematic of the kringle IV polypeptide chain. Amino acid positions mutated by Lp(a)-reducing missense variants are indicated in shades of red; cysteine residues forming disulfide bonds are highlighted in blue. **(E)** (top) KIV amino acid consensus sequence. (middle) Number of Lp(a)-reducing missense variants modifying each amino acid position (using the color scale in panel **D**). KIV domains 1–3 are merged because variants in KIV-1 exon 2 and KIV-3 exon 1 cannot be distinguished from KIV-2 variants. (bottom) Fraction of missense variants modifying each amino acid position that had PrimateAI-3D score *≥*0.5.

The allelic series of 95 missense variants in *LPA* that associated with greatly reduced Lp(a) levels implicated portions of the apo(a) amino acid sequence that are particularly sensitive to mutations (Fig. 5D,E). Most of the apo(a) sequence consists of kringle-IV repeats, including both the length-polymorphic KIV-2 repeat and nine other highly homologous sequences (KIV-1 to KIV-10). The 114 amino acid (aa) positions in the consensus repeat sequence (from EMBOSS Cons^63^) were altered by 329 missense variants included in our analyses, among which 78 variants were identified as likely to strongly reduce Lp(a). All 78 of these variants modified amino acids within kringle domains (78 aa); none affected the unstructured interkringle peptide chain (36 aa; Fig. 5D). Moreover, the specific amino acid positions implicated within kringle domains broadly matched computational predictions from PrimateAI-3D^64^ (Fig. 5E). These included several positions at or adjacent to conserved cysteine residues involved in disulfide bonds necessary for kringle structure; two mutations of such residues are known to encode apo(a) isoforms that cannot be secreted^23^. Lp(a)-reducing effect sizes of missense variants strongly correlated with computationally predicted variant effect scores (*R*=–0.58 between *β* and PrimateAI-3D score; Fig. 5C), providing further support for the robustness of these associations.

Together, these results provide a uniquely detailed inventory of how rare *cis* variation at a locus shapes population variation of a human phenotype, with genetic association data and machine learning-based variant effect prediction converging on a coherent picture of which portions of the *LPA* gene are key to its function.

### Changing selective pressure at *LPA* over the past million years

These data about the sensitivity of *LPA* to missense mutations, the allele frequencies of genetic determinants of Lp(a) levels, and the mutational properties of the KIV-2 repeat also provided insights into the trajectory of selection on *LPA* alleles over human evolutionary history. The high conservation of the *LPA* coding sequence across primates indicates that selective pressure for functional *LPA* has been present throughout most of primate evolution^6, 65^. However, the existence of common loss-of-function polymorphisms in humans and other primates^15, 66^ implies that *LPA* is no longer under selective constraint. Our data on the number of coding variants that damage *LPA* suggest that this cessation of selection for functional *LPA* occurred *<*1 million years ago: we estimate that *LPA* alleles are vulnerable to *>*8,400 damaging single nucleotide variants (Methods), such that at a de novo mutation rate of 1.2*×*10^−8^ per base pair per generation^67^, *LPA* function should be lost in the time scale of 30,000 generations in the absence of selection. In African populations, most *LPA* alleles are still functional, suggesting that selective constraint persisted until *<*1 million years ago.

Interestingly, in non-African populations, the high frequencies of *LPA* alleles that produce little or no Lp(a) provide some evidence that selection may even have reversed (in favor of null or hypomorphic alleles) in these populations in the past 50,000 years. Data from an ancient DNA-based analysis of directional selection suggest that SNPs at *LPA* associated with reduced Lp(a) levels have risen in frequency in European populations over the past 10,000 years^68^. The increased frequencies of alleles with longer KIV-2 repeats in non-African populations^69^ (Supplementary Table 2) despite the tendency of longer alleles to contract rather than expand (Fig. 2H) also suggest the possibility of recent selective pressure favoring longer alleles that produce less Lp(a). Selective pressure to reduce Lp(a) could reflect diminishing evolutionary benefits of Lp(a)^5, 6^ that have become outweighed by its atherogenic effects; future, larger ancient DNA data sets will help evaluate this hypothesis.

## Discussion

This work elucidates the genetic architecture of Lp(a) concentration across global populations, demonstrating how Lp(a) can be accurately predicted in multiple ancestries from allele-specific genotypes of KIV-2 length and functional sequence variants in *LPA*. Until measuring Lp(a) (at least once) becomes routine in clinical care^4^, genetic prediction could opportunistically identify people who would potentially benefit from future trials of Lp(a)-lowering therapies^70^, which may need to focus on primary prevention of cardiovascular disease^71^. The polygenic influences on Lp(a) identified here (such as the *APOE* L167del variant with opposite effects on Lp(a) and LDL) could also provide insights into Lp(a) metabolism, perhaps informing further therapeutic development^72^.

Lp(a) and the *LPA* locus also offer unique windows into genetics more broadly. Lp(a) is an extraordinary phenotype that is widely variable, nearly completely heritable, mostly monogenic, and accurately measured in hundreds of thousands of biobank participants. These properties enabled detailed genetic dissection of heritability at the *LPA* locus, yielding insights into how *cis*-epistasis and structural variation within a locus can impact polygenic score transferability—and showing how even for a mostly-monogenic trait, adjusting for genetic effects at a primary locus can reveal subtler, polygenic effects throughout the genome, consistent with the omnigenic model^73^. Studying abundant length polymorphism of the KIV-2 repeat using population-genetic approaches also yielded potentially generalizable insights into macrosatellite repeat mutation: the high mutability of the KIV-2 repeat appears to be driven by intra-allelic sister chromatid exchanges, suggesting that such mutations probably also occur somatically and that germline mutations are likely to be paternally-biased, increasing in frequency with paternal age^74^. Future data sets will enable further testing of these hypotheses for KIV-2 and other large multi-allelic CNVs.

Our study also had several limitations. Our estimates of KIV-2 allele lengths were quite accurate (RMSE=0.4) but still inexact, limiting the resolution of our analyses of allele-specific KIV-2 expansion and contraction rates and probably causing slight underestimation of the accuracy of our Lp(a) prediction model. Our data set also had limited representation of non-European ancestries, reflecting the composition of the UKB cohort (as Lp(a) measurements were largely unavailable and used heterogeneous measurement methods in AoU); as such, ancestry-specific Lp(a)-modifying variants surely remain to be discovered. Finally, while the full Lp(a) prediction model we developed is simple to describe, it is difficult to deploy, as it requires accurate, haplotype-resolved genotypes of KIV-2 repeat lengths together with SNP and indel variants in and near the repeat (though a possible workaround is to impute Lp(a) from SNP haplotypes^75, 76^). We anticipate that these challenges will be alleviated by future data sets with increased genetic diversity and new data types (e.g., ultra-high molecular weight sequencing or long-read sequencing combined with new assembly methods) together with continued development of scalable pipelines for short- and long-read analysis^77, 78^, enabling further insights into the genetics and evolutionary history of Lp(a).

## Methods

### Ethics

This research complies with all relevant ethical regulations. The study protocol (NHSR-4981) was determined to be not human subject research by the Broad Institute Office of Research Subject Protection as all data analyzed were previously collected and de-identified.

### UKB genomic data

UKB is a prospective cohort of volunteer participants aged 40–70 years at recruitment^79^. Short-read WGS data were previously generated from blood-derived DNA for 490,414 UKB participants using Illumina NovaSeq 6000 machines to an average coverage of 32.5*×*, and DRAGEN 3.7.8 was used to align 151-bp paired-end reads to GRCh38^33^.

We assigned genetic ancestries to UKB participants by using Euclidean distance^80^ to cluster them according to their first 20 genetic principal components (PCs) as previously described^41^ using PCs computed from SNP-array genotypes^79^. Of the 488,120 participants analyzed, 458,085 were assigned European genetic ancestry (EUR), 9,757 were assigned South Asian genetic ancestry (SAS), 7,848 were assigned African genetic ancestry (AFR), 1,488 were assigned East Asian genetic ancestry (EAS), and 10,942 were not assigned any genetic ancestry.

### Estimating diploid KIV-2 length from WGS read-depth

For each UKB WGS cram file, we computed a GC-adjusted estimate of diploid KIV-2 length (summed across the individual’s two alleles) by counting reads with alignment start positions in the region chr6:160611533–160646719 (GRCh38), adjusting for the sample’s GC profile in a manner following Genome STRiP v2^81^, as recently described^82^. Correcting for GC bias only partially accounted for technical variation in WGS read-depth, so to further denoise these diploid KIV-2 length estimates, we additionally computed “genomic wave” PCs capturing axes of technical bias in genome-wide read-depth profiles^82^; we then estimated each sample’s genomic wave pattern and adjusted its diploid KIV-2 length estimate according to the predicted amount of technical bias at KIV-2 (Supplementary Note 2).

### PSV-based KIV-2 phasing using haplogroup-informative PSVs

The DRAGEN KIV-2 CN caller introduced the idea of partitioning diploid KIV-2 copy number estimates into allele-specific length estimates using PSVs (i.e., SNPs within the KIV-2 region) with the property that nearly all KIV-2 repeat alleles are either homogeneous for the REF allele (throughout the repeat region) or homogeneous for the ALT allele^32^. Recombined alleles comprising hybrids of these haplogroups are extremely rare in the population (e.g., at most 2 out 5,008 alleles in 1KGP3; Supplementary Note 1). As such, individuals heterozygous for the PSV alleles can be assumed with high likelihood to carry one allele from each of the two KIV-2 haplogroups, allowing diploid KIV-2 length to be partitioned according to the proportion of WGS reads carrying REF vs. ALT alleles (across all PSV sites) in such individuals^32^.

Once diploid KIV-2 length has been accurately estimated, the key factor that limits the accuracy of this PSV-based partitioning approach is the stochastic sampling of allele-specific reads: only a fraction of the DNA fragments sampled from the KIV-2 region overlap PSVs, such that noise in the binomial sampling fraction introduces error in allele-specific length estimates. Another issue is that not all haplogroup-informative PSVs are perfectly homogeneous across KIV-2 alleles: in particular, one of the two PSVs used by the DRAGEN caller (represented by chr6:160613786:T*>*G) has the property that the ancestral PSV allele (T, the REF allele in GRCh38) is occasionally found in the first repeat unit of KIV-2 alleles belonging to the derived haplogroup (Fig. 1D and Supplementary Fig. 2). In carriers of such alleles, this “exception” to the PSV pattern creates a small bias in the allele-specific KIV-2 length estimates obtained by partitioning diploid KIV-2 copy number according to REF vs. ALT proportions.

To handle both of these issues, we incorporated a population-genetic approach into the PSV-based strategy. We first identified 3 of the 5 haplogroup-informative PSVs as being prone to common variation in the number of repeat units in which KIV-2 alleles from the derived haplogroup carried the reference PSV allele (Supplementary Fig. 2; a fourth PSV always exhibited exactly one such exception). KIV-2 alleles from the ancestral haplogroup contained almost no exceptions (Supplementary Fig. 2). Because the exceptions in KIV-2 alleles from the derived haplogroup nearly always occurred in the mutationally stable repeat units nearest to the flanks of the KIV-2 region (Fig. 1D), they were easy to impute from SNP-haplotypes. We performed imputation by, for each PSV, (1) genotyping PSV exception counts (i.e., the copy number of the reference PSV allele) in individuals homozygous for the derived (ALT) KIV-2 haplogroup; (2) phasing the exception copy numbers onto SNP-haplotypes^26^; and (3) imputing exception copy numbers into derived alleles of heterozygous individuals (Supplementary Fig. 3). We then performed PSV-based phasing adjusting for the imputed exception counts. Details of this procedure are described in Supplementary Note 3.

### SNP-based KIV-2 phasing using haplotype sharing

Because the PSV-based KIV-2 length estimation approach is only applicable to approximately half of the individuals in a cohort (namely, those heterozygous for the ancestral and derived haplogroups), we also computed allele-specific KIV-2 length estimates using our previous fully-population-genetic approach^26^. This approach, which relies on sharing of long (megabase-scale) SNP-haplotypes among individuals in a cohort who inherited the same allele, could obtain very accurate estimates of KIV-2 lengths for alleles shared by multiple closely or distantly related individuals. As such, SNP-based phasing was useful for maximizing sample size when analyzing effects of *LPA* variation on Lp(a) levels in EUR-ancestry UKB participants (even though PSV-based phasing was essential for studying non-European alleles and KIV-2 length mutation).

We used the same overall approach to SNP-based KIV-2 copy-number phasing that we previously described^26^. Starting with a phased SNP-array haplotype scaffold that we previously generated^83^, for each haplotype, we computed its longest identity-by-state (IBS) matches (of SNPs flanking KIV-2) to other SNP haplotypes in the UKB cohort^26^. We then phased diploid KIV-2 copy number estimates using the same probabilistic model of haplotype sharing and copy-number estimation error as before^26^. We optimized the parameters of the model using a slightly different approach in which we optimized imputation accuracy (which was easy to benchmark in a held-out sample set) rather than optimizing phasing accuracy, which we had previously done using a more complex procedure involving data from related individuals^26^.

### Genotyping coding and splice-region paralogous sequence variation at KIV-2

We genotyped paralogous sequence variants (PSVs) within the coding and splice regions of the KIV-2 repeat by analyzing WGS read alignments to GRCh38 previously generated using DRAGEN. For each sample, we ran samtools^84^ mpileup (options --excl-flags 772 --min-BQ 30 --no-BAQ --count-orphans) to count the number of reads supporting each allele (including indels) at each position within 20 bp of an *LPA* exon in the repeat region (chr6:160611560-160646781), which contains KIV-2 as well as parts of KIV-1 and KIV-3. We summed read counts of each observed base or indel across paralogous positions of repeat units in GRCh38, thus aggregating together variation throughout the variable number tandem repeat (VNTR) for PSV analysis. This approach did not attempt to disambiguate variants within the KIV-2 region proper from those in KIV-1 or KIV-3 (unlike ref.^77^).

At each profiled position, we estimated the diploid copy number (“dipCN”) of each observed PSV allele, summed across repeat units within both of an individual’s two repeat alleles, using the formula:

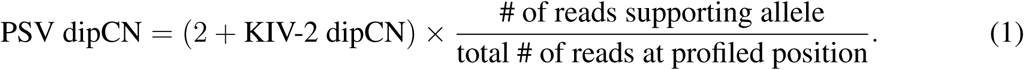

The addition of 2 repeat units to KIV-2 dipCN accounts for the overlap between the profiled VNTR region and the last exon of KIV-1 and the first exon of KIV-3 (which together contribute one repeat unit worth of exonic reads per allele, amounting to 2 additional repeat units per diploid genome). PSV dipCN estimates for alleles with fewer than 5 supporting reads in a given sample were set to 0. Variants whose mean dipCN across carriers (i.e., individuals with *≥*5 reads of support) was less than 0.75 were flagged as potential technical artifacts and excluded from further analysis.

We adopted a naming convention for PSVs with the form KIV2.EXON:POS*>*NT, where:

- EXON=1 or 2 is the index of the exon of the KIV-2 repeat that the variant lies in or near
- POS is the position relative to the start of the exon (in the direction of transcription, i.e., the minus strand of GRCh38):

**–** POS of “–X” indicates X bp before the start of the exon (i.e., to the right in GRCh38 coordinates)
**–** POS of “+X” indicates X bp after the end of the exon (i.e., to the left)
**–** POS of “X” indicates X bp into the exon (i.e., counting leftward from the rightmost base pair of the exon)
- NT is the nucleotide or sequence of nucleotides on the plus strand (opposite transcription) in the alternate allele (using as reference the GRCh38 sequence for the first two exons within the VNTR and their flanks: chr6:160644827–160645026 and chr6:160646194–160646415).

### Construction of SNP and KIV-2 PSV haplotypes at *LPA*

We generated a data set of phased *LPA* SNP+indel and PSV haplotypes as follows.

1. **Extract SNP and indel variants at *LPA* genotyped from WGS.** We used bcftools^85^ to extract variants within *∼*100 kb of *LPA* (chr6:160.42–160.78 Mb) from GraphTyper population level genotype call VCFs supplied by UKB^33^, filtering positions with more than 10 alleles, variants with fewer than 5 carriers, and variants with alternate allele “*”, and setting genotype calls with depth *<*10 or genotype quality *<*20 to missing. We excluded all GraphTyper variant calls within the KIV-2 region (chr6:160611533–160646869).
2. **Phase common WGS variants at *LPA***. We ran SHAPEIT5^86^ (v5.1.1, phase common with --region chr6:157490000-162500000) to phase the common (MAF*>*0.001) GraphTyper SNP and indel variants onto a scaffold of haplotypes previously generated from SNP-array genotypes^83^. We excluded variants with AAScore*<*0.8 or missingness*>*10%.
3. **Phase rare WGS variants and PSVs at *LPA***. We ran SHAPEIT5 (v5.1.1, phase rare) to phase rare PSVs in the KIV-2 VNTR (present in *≤*1% of alleles) and rare GraphTyper variants (MAF*≤*0.001, excluding variants with missingness *>*50%) onto the scaffold generated in step 2 above (phase rare with --region chr6:157490000-162500000 --scaffold-region chr6:157490000-162500000). For each genotyped rare PSV, we assigned each individual with *≥*5 reads of support a heterozygous genotype (“0/1”). Accuracy of haplotype phase obtained from phase rare is not influenced by the inclusion of poorly genotyped variants (e.g., SNPs with missingness up to *∼*50%), so we prioritized inclusivity in this step with the understanding that some variants may be low-quality.
4. **Phase common PSVs at *LPA***. We phased common PSVs using SNP haplotype sharing among UKB participants, using the same minor modifications to our previous VNTR phasing algorithm^26^ described above. This method takes as input dipCN estimates (which need not be integers and can exceed 2) and phases them into haploid copy number estimates.

### Variant annotation

We annotated variant consequences and locations within the *LPA* transcript using Variant Effect Predictor (VEP)^87^ v112.0. We scored potential impacts of variants on *LPA* splicing using SpliceAI^62^ v1.3.1, with option -D 4999. We obtained pre-computed PrimateAI-3D^64^ missense pathogenicity scores via application at https://github.com/Illumina/PrimateAI-3D.

### Lipoprotein(a) measurements in UKB

We obtained Lp(a) measurements for 456,283 UKB participants (including values outside the 3.8–189 nmol/L reportable range, available from Return 2321). Repeated measurements from a second visit to a UKB assessment center were available for 16,233 individuals. Measurements reported as “*>*567” were replaced with the value 657.4 nmol/L, the average of all reported specific values larger than 567 nmol/L. Measurements were averaged for individuals with multiple measurements.

To obtain a set of unrelated UKB participants, we iteratively removed one individual from each pair of individuals related at second degree or closer (kinship coefficient *>*0.0884 as computed by UKB^79^). We prioritized keeping individuals with an Lp(a) measurement, and otherwise chose a random individual among each related pair for removal.

The resulting data set consisted of 411,680 unrelated UKB participants who had a reported Lp(a) measurement, were assigned a genetic ancestry, and had SNP, indel and PSV haplotypes as well as phased KIV-2 allele lengths (from SNP-based phasing, and for approximately half of the cohort, also from PSV-based phasing).

### Predicting Lp(a) from genetic variation at the *LPA* locus

We generated Lp(a) predictions from phased *LPA* haplotypes using an optimized version of the model we described in our previous work^26^. Here, we fine-tuned the model by leveraging our larger data set with higher-accuracy KIV-2 allele lengths (n*∼*500,000 WGS-derived estimates vs. n*∼*50,000 WES-derived estimates in our previous work) and the availability of Lp(a) measurement data outside the “reportable range” of 3.8–189 nmol/L (which had not been available for our previous analysis).

#### Model definition

As in our previous work^26^, we modeled an individual’s Lp(a) concentration as the sum of contributions from the paternally- and maternally-inherited *LPA* alleles. Each allele’s contribution is a function of the allele’s phased KIV-2 length and SNP+indel haplotype, comprised of fine-mapped variants likely to causally modulate Lp(a) production:

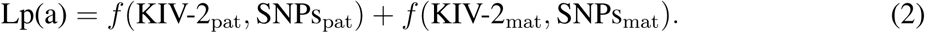

Here and below, we have written “SNPs” for brevity, but the set of fine-mapped variants (23 variants in our previous model, which we augmented here with 4 additional null-Lp(a) variants common in non-EUR ancestries) includes one indel among several PSVs within the KIV-2 region.

The contribution of each *LPA* haplotype is the product of a baseline contribution, determined by the allele’s KIV-2 repeat count, and a SNP haplotype effect:

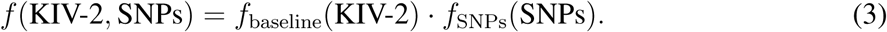

We previously chose *f* _baseline_ from a small parametric family to minimize model parameters and avoid overfitting^26^. Here, given the abundant sample size (tenfold larger than before), we made no assumptions about the functional form of *f* _baseline_, allowing it to be an arbitrary function on the possible integer KIV-2 copy numbers (cropping very long alleles to 28 repeat units, beyond which alleles produced negligible amounts of Lp(a): *≤*0.5 nmol/L according to our model). As in our previous work, each fine-mapped SNP used in the model was assumed to modify the amount of Lp(a) produced by haplotypes carrying the minor allele (defined according to EUR AF; Supplementary Table 3) by a fixed multiplicative factor:

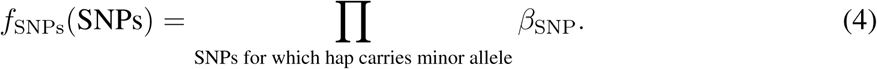

We classified the 23 SNPs carried over from our previous model into three categories according to their previously-estimated multiplicative effect sizes^26^ (which we subsequently fine-tuned as described below): null (*β* = 0), large-effect (0 *< β ≤* 0.5), and small-effect (*β >* 0.5). The parameters of our model thus corresponded to the baseline contribution of each integer KIV-2 copy number in {>1, 2, …, 28}> (Supplementary Table 2), the multiplicative effects of the large-effect SNPs (including separate multiplicative effect size estimates for 1 or 2+ copies of the two common cryptic splice PSVs), and the multiplicative effects of the small-effect SNPs (Supplementary Table 3). We set *β* = 0 for SNPs that appeared to produce null alleles.

#### Model-fitting

We fit the model parameters using data from unrelated UKB participants of European genetic ancestry. For model-fitting, we used KIV-2 allele lengths estimated for these individuals using our SNP-based phasing approach^26^ to maximize sample size. We adjusted raw Lp(a) measurements by the constant additive offset –1.51 nmol/L so that the average adjusted Lp(a) among individuals homozygous for the canonical splice variant rs41272114*>*T (which produces a common null allele^17^) was 0. We then performed model-fitting in two steps:

##### 1. Fit the baseline function *f* _baseline_ and the effect sizes of small-effect SNPs

To fit *f* _baseline_ and the effect sizes of the three small-effect SNPs (rs3124784*>*A, rs1853021:G, and rs1800769*>*T)^26, 88, 89^, we restricted analysis to individuals who did not carry any large-effect SNPs. If we hold the SNP effects fixed, then equation (2) is a linear function of *f* _baseline_(*k*) for *k ∈ {*1, 2*, …* 28}>. Likewise, equation (2) is a linear function of *β_s_* if we fix *f* _baseline_ and further restrict to individuals each of whose haplotypes carries at most one small-effect SNP *s*. Proceeding iteratively, starting from initial effect estimates *β_s_* = 1 for small-effect SNPs, we alternately generated least-squares estimates for *f* _baseline_(*k*) fixing *β_s_* and least-squares estimates for *β_s_* holding *f* _baseline_(*k*) fixed. We ran 15 iterations of this estimation procedure, at which point convergence had occurred.

While fitting *f* _baseline_(*k*) fixing *β_s_*, we constructed the design matrix (columns corresponding to integer KIV-2 copy numbers in {>1,2,…,28}>, rows corresponding to UKB participants) using the fractional part of KIV-2 allele length estimates to distribute contributions of individual alleles across matrix columns. A sample with an allele length estimate *k*_est_ had contributions to columns corresponding to *[k*_est_*♩* and *[k*_est_*♩* + 1 with weights 1*−*frac(*k*_est_) and frac(*k*_est_), where frac(*k*_est_) = *k*_est_*−[k*_est_*♩*. While fitting *β_s_* fixing *f* _baseline_(*k*), we excluded data from individuals with a long allele (KIV-2 length estimate *>*22.5 repeats) since the baseline contribution to Lp(a) for these haplotypes is small (*<*10 nmol/L; Supplementary Table 2).

To account for heteroskedasticity in Lp(a) measurements (which had higher variance for individuals with higher Lp(a)), we used inverse-variance weighted least squares when fitting *f* _baseline_(*k*) and *β_s_*. We used the following procedure to estimate the noise in each individual’s Lp(a) measurement and thus derive weights. At the end of each iteration, we used the current parameter estimates to generate an interim genetic prediction of Lp(a) for each UKB participant. We then used ordinary least squares to fit a model that estimated the squared error of measured Lp(a) using a quadratic function of predicted Lp(a):

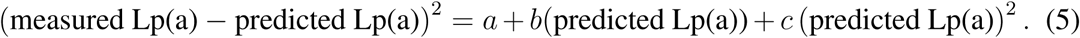

In the next iteration of parameter fitting, we weighted each row by the inverse of the predicted squared error, cropping the predicted squared error from below at 100. In the initial round of fitting, weights were initialized to 1.

##### 2. Fit effect sizes for large-effect SNPs

We estimated effect sizes for the large-effect SNPs by minimizing squared prediction error using linear regression. We first fit parameters for the two common large-effect SNPs (EUR AF*>*10%), both of which are PSVs (KIV2.2:182*>*T and KIV2.2:–11*>*T, i.e., G4925A and G4733A^21, 25^). We restricted to haplotypes with short and medium-length KIV-2 alleles (KIV-2 length estimate *<*19.5 repeats) that were opposite a haplotype carrying a null allele. We also required haplotypes to carry at most one of the two large-effect PSVs and none of the rarer large-effect SNPs. Since both common large-effect variants are PSVs, individual haplotypes sometimes carry multiple copies. We estimated separate effect sizes for haplotypes carrying exactly one copy (phased PSV copy number estimate in the range 0.5–1.5) and those carrying more than one copy (phased copy number estimate *>*1.5).

Finally, we fit parameters for the lower-frequency large-effect SNPs (EUR AF*≤*2%). We restricted to haplotypes with KIV-2 length *<*19.5 repeat units that carried exactly one of these SNPs. We further restricted to individuals whose opposite haplotype did not carry a rare large-effect SNP and was predicted to produce at most 50 nmol/L of Lp(a).

### Augmenting Lp(a) prediction with ancestry-specific SNPs from non-EUR fine-mapping

We ran stepwise conditional linear regression analyses separately in each non-European ancestry represented in UKB (AFR, EAS, SAS). At each stage of analysis, we tested the phenotype

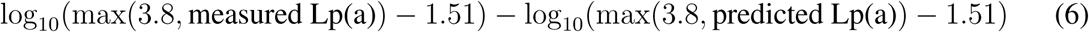

for association with all variants in the region chr6:160.42–160.78 Mb (extracted as described above). In the above formula, measured Lp(a) is on the “raw” scale with a mean of 1.51 nmol/L in null-allele homozygotes, and predicted Lp(a) is on the same scale via addition of a constant 1.51 nmol/L offset to equation (2).

We ran linear regression using BOLT-LMM^90, 91^ v2.5 with assessment center, genotyping array, sex, age, age^2^, and 20 genetic PCs as covariates, restricting analyses to unrelated individuals. We used Lp(a) predictions generated using PSV-based phased KIV-2 lengths for individuals with PSV-based phasing; for other individuals, we used predictions generated using SNP-based phased KIV-2 lengths. In the first round of analysis, Lp(a) predictions were generated using KIV-2 length and the 23 fine-mapped variants identified in our previous study^26^, with model parameters re-fit as described above. In subsequent rounds of analysis, Lp(a) predictions also included variants implicated in earlier rounds. Each such variant was categorized as a null allele by visual inspection of the distribution of measured Lp(a) in carriers (adjusted for the contribution of the opposite haplotype).

These analyses identified four protein-altering variants to add to the model, each of which was among the top associated variants in a round of analysis in a particular ancestry (Supplementary Fig. 5). In the first round of analysis in EAS, the strongest association involved the canonical splice acceptor variant KIV2.1:–1*>*T (*P*=2.7*×*10^−39^ vs. 3.7*×*10^−37^ for the next-strongest association). In the first round of analysis in AFR, the four strongest associations involved SNPs in LD (pairwise *R*^2^*>*0.88; *P*=5.6*×*10^−91^–4.6*×*10^−88^ vs. 4.1*×*10^−87^ for the next-strongest association), two of which (KIV2.1:93*>*A and chr6:160532622*>*G) are missense variants. In the second round of analysis, after including KIV2.1:93*>*A in the Lp(a) model, the strongest association involved the splice acceptor variant chr6:160578706*>*C (*P*=3.8*×*10^−62^ vs. 4.9*×*10^−49^ for the next-strongest association). In the third round of analysis, the strongest association involved the missense variant KIV2.1:64*>*T (*P*=1.4*×*10^−24^ vs. 7.8*×*10^−24^ for the next-strongest association). In the first round of analysis in SAS, the top variants included several plausibly-causal coding variants of large effect (stop-gained KIV2.2:108*>*C *P*=1.6*×*10^−20^; missense chr6:160532580*>*C *P*=5.7*×*10^−18^; missense KIV2.1:63*>*G *P*=1.4*×*10^−17^; stop-gained KIV2.1:9*>*A *P*=4.2*×*10^−14^). However, each of these variants is rare (SAS AF*<*0.2%) and including these variants in the Lp(a) model had a minor impact on prediction accuracy, so we did not add these variants to the model. The final Lp(a) model we used for assessing prediction accuracy included KIV-2 length, the 23 modifier variants from our previous model^26^, and the four additional variants KIV2.1:–1*>*T, KIV2.1:93*>*A, chr6:160578706*>*C, and KIV2.1:64*>*T (Supplementary Table 3).

### Comparison with published polygenic scores for Lp(a)

We compared the performance of our Lp(a) model with published polygenic scores from the PGS Catalog^92^ and our previous work^26^. We restricted analysis to unrelated UKB participants with PSV-phased KIV-2 allele lengths. We obtained SNP weights for each PGS for the “lipoprotein A measurement” trait (identifier EFO 0006925), excluding PGS’s derived from phenome-wide analyses (which typically do not account for the subtleties of the Lp(a) phenotype in UKB, in which commonly-observed measurements *<*3.8 nmol/L or *>*189 nmol/L were set to missing by default). We extracted SNP genotypes for PGS variants from the WGS-derived GraphTyper population level VCFs and computed scores for UKB participants using the reported effect weights. For PGS000752, we omitted 51 of 11,446 model variants that were either not in the UKB VCF or very rare in UKB (MAF*<*0.001); these exclusions did not affect the accuracy of the model, which had an accuracy in UKB individuals of EUR ancestry equal to the reported accuracy (*R*=0.52)^48^. We additionally compared our model to our previously published model using two different versions of *LPA* haplotypes: (1) those we previously derived from analysis of the initial release of WES data (n*∼*50,000)^26^, and (2) those we generated in this study from WGS data on the entire UKB cohort, using PSV-based phasing for KIV-2 length estimation.

### GWAS and rare variant association analyses of Lp(a)

We performed GWAS and exome-wide association analyses of inverse-normal transformed *LPA*-adjusted Lp(a) in UKB EUR-ancestry participants using BOLT-LMM (Supplementary Note 10). We designed a separate likelihood ratio test for effects of rare coding, splice region, and promoter variants in *LPA* (Supplementary Note 11).

### Estimating sensitivity of Lp(a) to coding variation at *LPA*

We estimated the rate at which *LPA* haplotypes mutate to acquire a LoF or near-LoF mutation by analyzing the numbers and fractions of such variants among the rare coding variants considered in our rare variant association analysis at *LPA*. For typical *LPA* alleles (*≥*10 KIV-2 repeats), *>*8,400 of the possible missense mutations are assigned a PrimateAI-3D score *>*0.5. Among variants in our association analysis with tested allele count *≥*10, *∼*70% of PrimateAI-3D *>*0.5 missense variants associated with *>*50% reduction in Lp(a). Furthermore, missense variants accounted for *∼*70% of all large-effect rare variant associations we identified. Together, this suggests that a typical *LPA* allele is vulnerable to *>*8,400 single nucleotide mutations. At a de novo mutation rate of 1.2*×*10^−8^ per bp per generation^67^, corresponding to a *∼*0.4*×*10^−8^ chance (on average) for a particular single nucleotide mutation per generation, a damaging *LPA* mutation should thereby occur at a rate *>*8,400 *×* 0.4*×*10^−8^, i.e., every *<*30,000 generations.

## Data availability

Genomic and phenotypic data analyzed in this study are available by application to UK Biobank (http://www.ukbiobank.ac.uk/) and All of Us (https://allofus.nih.gov/). Full summary statistics from GWAS of *LPA*-adjusted Lp(a) in UKB will be made available at the NHGRI-EBI GWAS Catalog after completion of peer review. Individual-level *LPA* haplotypes and Lp(a) predictions in AoU v7 will be released in a community workspace, and individual-level *LPA* haplotypes and Lp(a) predictions in UKB will be returned when a mechanism for release on the UKB-RAP platform becomes available.

## Code availability

Code used to perform analyses reported in this study will be publicly released after completion of peer review.

## Acknowledgements

We thank P. Palamara and M. Levin for helpful discussions. This research was conducted using the UK Biobank Resource under application no. 40709. R.E.M. was supported by US NIH grant K25 HL150334; S.R. by a Swiss National Science Foundation Postdoc.Mobility fellowship (P500PB 211106); N.K. by US NIH fellowship F31 DE034283; R.E.H. and S.A.M. by US NIH grant R01 HG006855; M.L.A.H. by a US NIH fellowship F32 HL160061; D.T. by US NIH training grant T32 HG002295; and P.-R.L. by US NIH grants R56 HG012698 and R01 HG013110 and a Burroughs Wellcome Fund Career Award at the Scientific Interfaces. The funders had no role in study design, data collection and analysis, decision to publish or preparation of the manuscript. The content is solely the responsibility of the authors and does not necessarily represent the official views of the National Institutes of Health. Computational analyses were performed on the UKB Research Analysis Platform, the All of Us Researcher Workbench, and the O2 High Performance Compute Cluster supported by the Research Computing Group at Harvard Medical School. The All of Us Research Program is supported by the NIH, Office of the Director: Regional Medical Centers: 1 OT2 OD026549; 1 OT2 OD026554; 1 OT2 OD026557; 1 OT2 OD026556; 1 OT2 OD026550; 1 OT2 OD026552; 1 OT2 OD026553; 1 OT2 OD026548; 1 OT2 OD026551; 1 OT2 OD026555; IAA no. AOD 16037; Federally Qualified Health Centers: HHSN 263201600085U; Data and Research Center:5 U2C OD023196; Biobank: 1 U24 OD023121; The Participant Center: U24 OD023176; Participant Technology Systems Center: 1 U24 OD023163; Communications and Engagement: 3 OT2 OD023205; 3 OT2 OD023206; and Community Partners: 1 OT2 OD025277; 3 OT2 OD025315; 1 OT2 OD025337; and 1 OT2 OD025276. In addition, the All of Us Research Program would not be possible without the partnership of its participants.

## Author contributions

R.E.M. and P.-R.L. conceptualized the project, performed analyses, and drafted the manuscript. S.R., N.K., R.E.H., M.L.A.H., and D.T. assisted with analytical pipelines and interpretation of results. All authors reviewed and edited the manuscript.

