## Supplementary Information for "The genetic architecture of lipoprotein(a): simplicity, complexity, and mutability"

#### Contents

**Supplementary Notes** 2

**1 Haplogroup structure and recombination at the KIV-2 locus** 2

**2 Denoising WGS read-depth at KIV-2** 3

**3 PSV-based phasing of KIV-2 allele lengths** 5

**4 Benchmarking accuracy of PSV-based KIV-2 allele length estimates** 9

**5 Estimating copy-number mutation rate of the KIV-2 repeat** 12

**6 Factors limiting SNP-based prediction of Lp(a)** 16

**7 Estimating Lp(a) heritability attributable to the *LPA* locus** 17

**8 Genetic architecture of Lp(a) across ancestries** 19

**9 Lp(a) prediction and cardiovascular diseases in AoU** 20

**10 Genome-wide association analysis of Lp(a) adjusted for *LPA* variation** 22

**11 Testing rare coding, splice region, and promoter variants for association with Lp(a)** 24

**References** 28

**Supplementary Figures** 37

**Supplementary Tables** 46

### Supplementary Notes

#### 1 Haplogroup structure and recombination at the KIV-2 locus

**Genealogy of KIV-2 alleles.** We used RelateExtract to extract a coalescent tree at the KIV-2 locus (GRCh37 6:161071212) from an ancestral recombination graph previously inferred by Relate[35] for 302 modern human individuals (1000 Genomes CHB, GBR, YRI) and four archaic hominin individuals (the 1000G_sub_Nea data set generated by ref.[35]). We used newick-utils-1.6 (ref.[93]) to downsample modern human haplotypes by a factor of 10 for plotting in Fig. 1A. We used a generation time of 28 years as suggested in the Relate documentation.

**LD and recombination at the KIV-2 locus.** We computed linkage disequilibrium between rs6902102 (which is 7 kb downstream of the KIV-2 repeat and segregates with the deep split in KIV-2 haplotypes) and other common SNPs in the KIV-2 region (MAF>0.01 in the 1000 Genomes Phase 3 genotype call set; n=2,504) using plink2 (ref.[94]). We estimated the genetic map distance spanning the KIV-2 repeat by averaging across the 20 population-specific recombination maps previously generated for phased Omni genotypes for 1000 Genomes populations[37] using LDhat[36].

We downloaded the Halldorsson et al. 2019 data set containing genomic locations of 4,531,536 meiotic crossovers observed across 126,427 meioses[38]. Of these crossovers, 3,327,272 (73%) had been localized to regions of GRCh38 spanning <100kb, none of which spanned the KIV-2 repeat.

**Recombined KIV-2 repeat alleles.** We searched the 1KGP3 data set (n=2,504 individuals; 5,008 haplotypes) for evidence of KIV-2 alleles that might have recombined within the KIV-2 repeat (chr6:160611533–160646869). To do so, we identified two SNPs in each 1-kb flank of the KIV-2 repeat that were in high LD with rs6902102 (chr6:160610686:A:T and chr6:160611057:T:C on the left; chr6:160647032:G:C and chr6:160647144:T:G on the right in GRCh38 coordinates). We then searched for 1KGP3 participants who had consistent genotypes (i.e., the same number of ALT alleles) at the two left-flank SNPs and likewise had consistent genotypes at the two right-flank SNPs (providing a safeguard against genotyping error) but had discordant genotypes between the left and right flanks (suggesting the presence of a recombined allele, the left side of which matched one haplogroup and the right side of which matched the other). We identified exactly 2 individuals with this property, both of African ancestry: NA18917 from the YRI population and NA19437 from the LWK population.

Examining the Relate coalescent tree at KIV-2 showed that NA18917 and NA19437 shared a haplotype that formed a clade, indicating that these two individuals probably co-inherited an allele formed by a single, relatively recent recombination event, potentially within the KIV-2 repeat. We could not conclusively determine whether the recombination occurred in the KIV-2 repeat itself or in the few hundred base pairs immediately flanking the KIV-2 region.

**Haplogroup-informative paralogous sequence variants within KIV-2 alleles.** We searched for PSVs with near-perfect intra-allelic homogeneity using a combination of short-read WGS analyses (looking for common PSVs with high mean copy number—estimated by partitioning read-depth-derived repeat copy number according to REF and ALT counts at the PSV[95]—but near-zero copy number in a substantial fraction of individuals) and analyses of long-read data from HGSVC2[96] and assemblies from HGSVC3[39] (accession PRJEB76276). These approaches converged on five haplogroup-informative PSVs (including one dinucleotide variant; Supplementary Table 1) that were all in strong linkage with rs6902102.

We identified the locations of these and other PSVs within assembled KIV-2 alleles from HGSVC3 (Fig. 1D,E) using the following procedure. First, we created a 5544-bp consensus KIV-2 repeat unit from five repeat units within GRCh38 (with boundaries at 0-based BED positions 160617116, 160622662, 160628206, 160633752, 160639299, and 160644846) using MUSCLE[97] and EMBOSS Cons[63]. We then used BLASTN[98] to identify assembly segments matching this sequence, which we extracted using samtools[84]. Finally, we aligned each repeat unit extracted from the assembly to the consensus repeat unit using minimap2[99], and we used samtools mpileup to identify variants.

### 2 Denoising WGS read-depth at KIV-2

As outlined in Methods, after performing GC-bias correction, we estimated and corrected for remaining technical biases using a principal component analysis (PCA)-based approach similar to our previous work[82]. As in our previous work, our overall approach was to compute top "genomic wave" PCs that captured axes of technical bias in genome-wide read-depth profiles. We could then estimate each sample's coefficients along these PCs, thus obtaining an estimate of the sample's genomic wave pattern that we used to further calibrate its read-depth-derived estimate of diploid KIV-2 copy number.

To use this approach to denoise KIV-2 copy number estimates, we generated genome-wide GC-corrected, CNV-masked WGS read-depth profiles at $\sim$50-kb resolution for a set of reference samples (UKB participants of age $\leq$44 years with sample contamination $\leq$0.002 and no mosaic chromosomal alterations previously detected by SNP-array analyses) as previously described[82]. These read-depth profiles (and genomic wave patterns estimated from them) excluded copy-number-variable regions such as KIV-2; however, they could still be used to denoise read-depth at KIV-2 by normalizing against a sample's estimated genomic wave value in the region flanking KIV-2 (specifically, chr6:160.599–160.691 Mb, the bin containing KIV-2—which had been masked from the read-depth profiles—along with $\sim$50 kb of unmasked flanking sequence).

We used this approach to denoise diploid KIV-2 copy number estimates in our benchmarks of our pipeline on 1000G samples (Supplementary Note 4). However, using the estimate of a sample's genomic wave value in this flanking region to proxy for its genomic wave value within KIV-2 was suboptimal, so for UKB, we devised an enhanced denoising approach that directly estimated each sample's genomic wave value within KIV-2.

To do so, we augmented each sample's read-depth profile with one additional measurement that estimated technical bias in WGS read-depth in the KIV-2 region. To obtain this estimate, we needed not only (i) the observed GC-corrected read-depth in the KIV-2 region (computed as described in Methods) but also (ii) the sample's expected read-depth at KIV-2 based on its diploid KIV-2 copy number—estimated in an independent manner (i.e., not derived from the sample's WGS read-depth). Given that WES data were available for the majority of the UKB cohort, we obtained these independent estimates of KIV-2 copy number from denoised WES read-depth[95] (across homologous copies of KIV-2 exon 2, which were captured by the WES oligo lots used to sequence all but the initial 50K WES samples); we further refined these estimates using our SNP-based phasing approach[26]. These refined WES-based KIV-2 copy number estimates were sufficiently accurate (in UKB EUR-ancestry participants) to discern evenly spaced modes corresponding to integer copy numbers. This allowed us to calibrate the WES-based KIV-2 copy number estimates to integer modes, after which we subtracted $2 \times 0.65$ to account for the WES-based estimates measuring an extra repeat unit per haplotype (owing to counting the second exon of KIV-1 along with the second exons of the KIV-2 repeats) whereas our WGS read-depth measurements in the KIV-2 region measured an extra $\sim$0.35 repeat units per haplotype.

After augmenting the reference read-depth profiles with these estimates of technical bias at KIV-2 (i.e., ratios of (i) to (ii) above), we ran PCA on the EUR subset of the reference cohort using the same approach as before[82]. Finally, for each UKB WGS cram file, we estimated its genomic wave pattern by computing the coefficients of its $\sim$50-kb resolution read-depth profile along the top 40 PCs. The linear combination of the 40 PCs defined by these coefficients gave the estimated technical bias across the sample's genome-wide read-depth profile. Applying these coefficients to the components of the PCs at KIV-2 (i.e., the augmented coordinate of the read-depth profile vectors) provided an estimate of the extent to which WGS read-depth over/under-estimated the sample's KIV-2 copy number. We thus obtained our final estimate of the diploid KIV-2 copy number of each UKB participant by dividing GC-corrected KIV-2 read-depth by this PC-derived estimate of technical bias and subtracting $2 \times 0.36$ to remove the contribution of the extra partial repeat unit per haplotype that had been included in the WGS read-depth measurement.

We note that the increased accuracy afforded by this enhanced PCA-based denoising procedure was important primarily for our analyses of KIV-2 mutation (in which being able to distinguish mutations from measurement noise was crucial). The increased KIV-2 length accuracy was less essential for studying genetic effects on Lp(a); here, the enhanced denoising primarily improved the accuracy of our estimates of Lp(a) prediction accuracy, as measurement error of KIV-2 allele lengths leads to downward bias in this quantity. We also note that in the absence of orthogonal WES data, the enhanced PCA-based denoising approach described above could still be applied by instead using SNP-based imputation of WGS-derived KIV-2 copy numbers to obtain diploid KIV-2 copy number estimates independent of an individual's own WGS read-depth data.

### 3 PSV-based phasing of KIV-2 allele lengths

The approach that we used to estimate allele-specific KIV-2 repeat lengths using five haplogroup-informative PSVs (building on the DRAGEN KIV-2 CN caller[32]) is overviewed in Methods. This supplementary note describes details of our implementation.

**Estimating diploid copy numbers of haplogroup-informative PSVs.** For each UKB WGS cram file, we estimated the diploid copy number of each of the five haplogroup-informative PSVs (Supplementary Table 1) by partitioning diploid KIV-2 copy number (denoised as described in Supplementary Note 2) according to the relative counts of reads containing its REF allele vs. ALT allele. Because sequencing and read-mapping biases can create slight biases in the allelic depths observed for different alleles of a variant, we took two steps to minimize such biases.

First, to minimize REF-bias (introduced by soft-clipping of read alignments containing an alternate allele near the end of the read, causing the alignment to be truncated and the read not to contribute its ALT allele to the allelic depth count), we generated an "ALT contig" in which we replaced the REF allele in each GRCh38 position of each of the five PSVs (Supplementary Table 1) with the ALT allele of the PSV. Additionally, because one of the PSVs ("PSV 2222") is typically present as a dinucleotide variant, we also replaced the preceding base (G) with its ALT allele (T) at all of its GRCh38 locations. We created a REF+ALT reference FASTA file containing the original GRCh38 sequence chr6:160.6–160.65 Mb (50 kb containing the KIV-2 region and its flanks, extracted using bedtools[100]) along with the corresponding ALT contig.

To tabulate allelic depths using this augmented reference, we used samtools[84] to extract WGS reads that DRAGEN had aligned to chr6:160601000-160649000 (excluding SAM flags 772: unmapped, non-primary, QC-fail) and used bwa mem[101] to realign the extracted reads to our REF+ALT reference. We then used samtools mpileup --count-orphans --no-BAQ --excl-flags 772 --min-BQ 20 --output-QNAME to tabulate reads supporting each allele at each PSV position (restricting to base calls with quality $\geq$20). We post-processed the mpileup output to collate allelic depths across all positions of each PSV and to ensure that each read name (corresponding to a single DNA fragment) was counted at most once in the allelic depth tally of each of the five PSVs (preventing double-counting of fragments for which the forward and reverse read in a pair overlapped). Finally, we rounded any allelic depth $<$4 down to 0, as small nonzero allelic depths were likely to reflect sequencing errors.

The above procedure largely eliminated REF-bias, but we observed a small ($\sim$1%) residual bias in allele counts that appeared to be base-specific: i.e., certain bases generated slightly higher allelic depths (per repeat copy) than others. We quantified the mean allelic depth generated per copy of each PSV allele by comparing allelic depths of PSVs to diploid KIV-2 copy numbers in individuals fully homozygous for either the PSV REF allele or the PSV ALT allele. This analysis indicated that the slight residual bias could be corrected using a second, simple adjustment: empirically, C bases exhibited 1.009-fold upward-bias and G bases exhibited 1.015-fold upward-bias in allelic depths (per repeat copy) relative to A and T bases, so we divided C and G allelic depths of PSVs by these factors.

We thus obtained the following formulas estimating the number of repeat units (summed across an individual's two alleles) carrying the REF allele for the five PSVs of interest:

$$\text{PSV 706 G>C:} \qquad \text{REF dipCN} = (\text{KIV-2 dipCN} + 2) \times \frac{\text{AD}_\text{G}/1.015}{\sum \text{AD}/\text{bias}} \tag{7}$$

$$\text{PSV 1935 T>C:} \qquad \text{REF dipCN} = (\text{KIV-2 dipCN} + 2) \times \frac{\text{AD}_\text{T}}{\sum \text{AD}/\text{bias}} \tag{8}$$

$$\text{PSV 2222 T>G:} \qquad \text{REF dipCN} = (\text{KIV-2 dipCN}) \times \frac{\text{AD}_\text{T}}{\sum \text{AD}/\text{bias}} \tag{9}$$

$$\text{PSV 3190 C>G:} \qquad \text{REF dipCN} = (\text{KIV-2 dipCN}) \times \frac{\text{AD}_\text{C}/1.009}{\sum \text{AD}/\text{bias}} \tag{10}$$

$$\text{PSV 4929 A>G:} \qquad \text{REF dipCN} = (\text{KIV-2 dipCN}) \times \frac{\text{AD}_\text{A}}{\sum \text{AD}/\text{bias}} \tag{11}$$

where each denominator contains the total read-depth of the PSV (collated across repeat units in the reference) after correcting C and G allelic depths for their empirical biases. The PSVs are named according to their positions within the 5544-bp consensus KIV-2 repeat unit that we generated (Supplementary Note 1); the sets of GRCh38 coordinates corresponding to each PSV are indicated in Supplementary Table 1. The first two PSVs (706 and 1935) fall within the $\sim$35% of the KIV-2 repeat unit that is present in an additional copy on each allele (owing to the region of near-perfect homology extending beyond KIV-2 into the beginning of KIV-3 and the end of KIV-1); for this reason, the copy numbers of these PSVs are computed by partitioning KIV-2 dipCN + 2 rather than KIV-2 dipCN. These PSV copy number estimates include the potential contributions of the additional copy per allele.

**Phasing and imputing PSV exception counts using SNP haplotypes.** As noted in Methods, KIV-2 alleles from the ancestral haplogroup contained only the REF allele of all five haplogroup-informative PSVs (with negligibly rare exceptions), but KIV-2 alleles from the derived haplogroup were not fully homogeneous for the ALT allele of all PSVs (Supplementary Fig. 2): only PSV 3190 (represented by chr6:160614754) had this property. PSV 706 (represented by chr6:160612270) was the next-simplest, with derived-haplogroup KIV-2 alleles always containing exactly one repeat unit (the first) with the REF allele. The other three PSVs exhibited common variation in the number of repeat units in which derived-haplogroup KIV-2 alleles carried REF PSV alleles ("exceptions").

To enable accounting for these exceptions when using PSV copy numbers to phase individuals heterozygous for the two haplogroups, we used the following SNP-haplotype-based imputation strategy. First, we generated a reference panel consisting of UKB participants homozygous for the derived KIV-2 haplogroup. For each of these individuals, we estimated the diploid copy number of the REF allele of PSV 1935, 2222, and 4929 (represented by chr6:160646781, chr6:160613786, and chr6:160644231) as described above. This diploid REF copy number corresponded to the individual's total number of exceptions for the PSV, summed across the individual's two derived-haplogroup alleles. Because exception counts were low (with most alleles containing 0 or 1 exception for each PSV), these diploid REF copy numbers were estimated reasonably accurately from WGS read-depth (Supplementary Fig. 2). We then applied our previous SNP-based phasing and imputation approach[26] (with the minor modification to the parameter optimization routine described in Methods) to phase the continuous-valued exception count estimates in the hom-derived panel and impute exception count dosages for PSV 1935, 2222, and 4929 into het-ancestral/derived individuals (i.e., the individuals for whom we wished to perform PSV-based KIV-2 phasing).

In the above analyses, we defined KIV-2 haplogroup genotypes (hom-derived for the imputation panel, het-ancestral/derived for the target individuals) using four SNPs (two on each flank of the KIV-2 repeat) in near-perfect LD with PSV 3190: chr6:160602177, chr6:160604256, chr6:160649838, and chr6:160650180. We restricted the imputation panel to individuals who were hom-ALT for all four tag SNPs, and we restricted the target cohort to individuals who were heterozygous for all four tag SNPs. This guarded against genotyping error and ensured that rare recombined alleles (with part of the allele originating from one haplogroup and the other part from the other haplogroup)—for which PSV-based phasing would produce incorrect results—were excluded from the analysis.

**Filtering to individuals with high-confidence PSV data.** In addition to restricting the target sample set to UKB participants heterozygous for the four haplogroup-tagging SNPs listed above (n=223,397), we applied two additional filters to further increase the robustness of the sample set on which to perform PSV-based phasing of KIV-2 allele lengths. First, we dropped 468 individuals for which one or more of the five haplogroup-informative PSVs had evidence of a third base (supported by $\geq$7 reads) beyond the expected REF and ALT bases. Such an observation indicated the presence of a mutation at the PSV position in one or more repeat copies, which would introduce error when using PSV REF vs. ALT allelic depths to partition KIV-2 diploid copy number into allele-specific components.

Second, we dropped 6,993 individuals with shorter SNP-haplotype matches to haplotypes in the imputation panel. Here, the rationale was to restrict the target set to individuals with very accurately imputed PSV exception count dosages. We implemented this filter by recording, for each target haplotype, a metric reflecting the lengths (in cM coordinates) of IBS matches to the haplotypes from the imputation panel that had been used to impute the target haplotype's PSV exception counts. Specifically, for each such IBS match to the panel, we computed its "IBS edge" distance as the shorter of the two distances from the KIV-2 repeat to the left and right edges of the IBS tract, and we computed the target haplotype's overall IBS edge distance as the weighted average across its IBS matches (using the same weights used during imputation[26]). Finally, we defined our filter by restricting to target individuals for whom one haplotype had IBS edge distance at least 0.1 cM longer than the other. This filter both enforced a minimum length of IBS matching and ensured that the identities of the target SNP-haplotypes carrying the ancestral-haplogroup vs. derived-haplogroup alleles were clear (which was necessary because our SNP haplotype scaffold[83] contained only SNP-array variants). For most target individuals, one haplotype (from the derived haplogroup) had a long IBS edge distance and the other haplotype (from the ancestral haplogroup) had near-zero IBS edge distance, as it did not match haplotypes from the hom-derived imputation panel.

**Estimating allele-specific KIV-2 repeat lengths.** For each remaining het-ancestral/derived target individual that satisfied the above filters, we computed the following estimates of the length of the individual's ancestral-haplogroup (REF) allele (one estimate from each of the five haplogroup-informative PSVs):

$$\text{PSV 706 G>C:} \quad \widehat{\text{KIV-2 CN}}_{\text{ancestral-hap allele}} = (\text{PSV 706 G dipCN}) - 1 - 1 \tag{12}$$

$$\begin{aligned}\text{PSV 1935 T>C:} \quad \widehat{\text{KIV-2 CN}}_{\text{ancestral-hap allele}} &= (\text{PSV 1935 T dipCN}) \\ &- (\text{imputed PSV 1935 T dosage on derived-hap allele}) - 1\end{aligned} \tag{13}$$

$$\begin{aligned}\text{PSV 2222 T>G:} \quad \widehat{\text{KIV-2 CN}}_{\text{ancestral-hap allele}} &= (\text{PSV 2222 T dipCN}) \\ &- (\text{imputed PSV 2222 T dosage on derived-hap allele})\end{aligned} \tag{14}$$

$$\text{PSV 3190 C>G:} \quad \widehat{\text{KIV-2 CN}}_{\text{ancestral-hap allele}} = (\text{PSV 3190 C dipCN}) \tag{15}$$

$$\begin{aligned}\text{PSV 4929 A>G:} \quad \widehat{\text{KIV-2 CN}}_{\text{ancestral-hap allele}} &= (\text{PSV 4929 A dipCN}) \\ &- (\text{imputed PSV 4929 A dosage on derived-hap allele})\end{aligned} \tag{16}$$

where the first terms in these equations are the PSV REF diploid copy numbers from equations (7)–(11). The second term of each equation subtracts off the number of PSV REF copies (i.e., exceptions) on the opposite (derived-haplogroup) allele: 1 copy for PSV 706, and the imputed numbers of copies for PSVs 1935, 2222, and 4929. Finally, an additional 1 copy is subtracted off in equations (12) and (13) because PSV 706 and 1935 occur in one additional non-KIV-2 repeat unit (KIV-1 or KIV-3) within the extra ~0.35 repeat unit of homology.

For most target individuals, we obtained the final estimate of the ancestral-haplogroup KIV-2 allele length by averaging across the five estimates from equations (12)–(16). For a small subset of individuals with a high imputed dosage of PSV 4929 exceptions (>2.5 copies, which was rare in UKB; bottom row of Supplementary Fig. 3), we excluded the estimate from PSV 4929 in equation (16) and used the average across the other four estimates, as the imputed exception count was less accurate in such cases (because the aberrant PSV alleles were not confined to the first or last repeat units of the KIV-2 repeat and thus were subject to increased copy-number mutation). Finally, we estimated the derived-haplogroup KIV-2 allele length as the diploid KIV-2 copy number minus the ancestral-haplogroup copy number.

#### 4 Benchmarking accuracy of PSV-based KIV-2 allele length estimates

We performed two sets of benchmarks of our PSV-based approach for estimating allele-specific KIV-2 repeat lengths: (1) benchmarks in UKB using IBD2 sibling pair concordance; and (2) benchmarks in 1000G using gold-standard Bionano optical genome mapping data (from ref.[32]). In the latter benchmarks, we also benchmarked allele length estimates previously generated using the DRAGEN KIV-2 CN caller[32] (which had been run on 1000G samples but not UKB) for comparison.

**UKB benchmarks using IBD2 sibling pairs.** We benchmarked the accuracy of allele-specific PSV-phased KIV-2 length estimates in UKB using sibling pairs IBD2 at *LPA* (based on at most three mismatching SNP-array genotypes within a 2-Mb window centered at KIV-2, computed using plink1.9 --genome). We restricted to sibling pairs each heterozygous for one ancestral and one derived-haplogroup allele. We determined that the PSV-based allele length estimates were well-calibrated based on their clustering into discrete modes at integer values (Fig. 2A,C). As such, the squared deviation between estimates of an allele's length in two siblings provided an estimate of twice the squared error of each estimate, i.e., RMSE $\approx$ RMSD$/\sqrt{2}$. This error estimate was expected to be slightly inflated by de novo mutations that occasionally caused true allele length discordances between IBD2 siblings, so we excluded the three highly discordant sibling pairs (Fig. 2A) and confirmed that the resulting error estimate (RMSE=0.4 repeat units) was consistent with an estimate based on the median absolute deviation (MAD), which is robust to outliers.

**1000G benchmarks using gold-standard Bionano data.** We also directly benchmarked our PSV-based KIV-2 length estimation approach (along with previously reported DRAGEN KIV-2 copy number estimates[32]) against gold-standard Bionano optical genome mapping data previously generated for a subset of 1000G samples (published in Supplementary Table 1 of ref.[32]).

We first verified that the Bionano data had a distribution consistent with these estimates being near-perfect. We focused on 87 samples with Bionano allele length estimates available for both alleles and indicated as high-confidence. These 174 allele length estimates were tightly clustered around integers, with a slight mean offset of +0.09 repeat units and a standard deviation of 0.12 repeat units around the offset. Based on this information, we concluded that the Bionano estimates had minimal error. In our 1000G benchmarks (Supplementary Fig. 4), we subtracted the estimated 0.09-repeat-unit offset from the Bionano length estimates published in ref.[32], but the choice of doing so versus using the estimates as-is had a negligible impact on the benchmarks.

Among the 87 samples with complete, high-confidence Bionano data, 85 samples were included in the 1000G high-coverage WGS data set[40] and could be used for benchmarking diploid copy number estimates produced by our pipeline and DRAGEN. Among these 85 samples, 55 samples were het-ancestral/derived and could be analyzed using PSV-based allele-specific copy number analysis. Because the haplotype correspondences between the Bionano data and the allele-specific length estimates obtained from WGS (by our analysis or DRAGEN) were unknown, we further restricted analysis to 49 of the 55 heterozygous samples for which the Bionano allele length estimates for the two haplotypes differed by $>2.5$ repeat units; this deviation was sufficiently large that the shorter WGS-derived length estimate could be assumed to correspond to the shorter of the two allele lengths in the Bionano data.

We ran our PSV-based KIV-2 length estimation method on the 1000G high-coverage WGS data set[40] (n=3,202) using alignments to GRCh38 from DRAGEN v4.0.3 that were available via the Registry of Open Data on AWS (`https://registry.opendata.aws/ilmn-dragen-1kgp/`). We performed read-depth profiling of each sample and computed PCs estimating genome-wide genomic wave patterns (at ~50-kb resolution) as previously described[82]. We then estimated each sample's diploid KIV-2 copy number by computing its GC-adjusted count of reads aligned to the KIV-2 region (as described in Methods) and dividing by its estimated genomic wave value in the chr6:160.599–160.691 Mb bin (i.e., we applied the basic PCA-based denoising approach described in Supplementary Note 2). Two samples had missing cram index files and were dropped from downstream analysis. To rescale these denoised KIV-2 read-depth estimates (which had been normalized such that a sample homozygous for the GRCh38 reference sequence should have a value of 1) to calibrated diploid copy number estimates, we multiplied by $2\times6.35$ (because GRCh38 contains 6 copies of the KIV-2 repeat along with the partial ~0.35-repeat worth of near-identical sequence in KIV-1 and KIV-3) and subtracted $2\times0.35$ to obtain estimates of diploid counts of full KIV-2 repeat units.

We then followed the procedure described in Supplementary Note 3 to (i) estimate diploid copy numbers of haplogroup-informative PSVs, (ii) phase PSV exception dosages in samples homozygous for the derived haplogroup (n=546) and impute them into het-ancestral/derived samples (n=1,504) using phased 1000G SNP-haplotypes[40] restricted to common SNPs on the UKB SNP-array, and (iii) estimate allele-specific KIV-2 repeat lengths by partitioning diploid copy number estimates according to the allelic depths of the five haplogroup-informative PSVs, adjusting for imputed exception dosages. We filtered 9 samples for which one or more of the five haplogroup-informative PSVs had evidence of a third base (supported by $\geq$7 reads) beyond the expected REF and ALT bases, leaving 1,495 samples with PSV-phased KIV-2 allele length estimates. These samples could alternatively have been retained by excluding allelic depth data from the PSV with the third base.

We benchmarked these WGS-based KIV-2 copy number estimates and previously published estimates generated by DRAGEN v4.2 (from Supplementary Table 2 of ref.[32]) against the gold-standard Bionano data, benchmarking the accuracy of both diploid copy number estimates (Supplementary Fig. 4A,B) and allele-specific copy number estimates (Supplementary Fig. 4C,D). The results of these benchmarks were as follows:

- For diploid KIV-2 copy number estimation, our read-depth denoising approach (which corrected for GC bias using a sample's genome-wide GC profile and then used PCA to adjust for its estimated genomic wave profile near KIV-2) produced considerably more accurate estimates than DRAGEN (which corrected for GC bias using 3,000 normalization regions of 2 kb each[32]): RMSE=0.73 copies for DRAGEN (Supplementary Fig. 4A) versus RMSE=0.49 copies from our method (Supplementary Fig. 4B). As expected, the DRAGEN benchmark was very consistent with the benchmark reported in ref.[32] (median difference in total copy number (vs. Bionano) reported to be $\sim$0.47 copies[32]); in our benchmark (which restricted to the subset of samples with Bionano calls indicated as high-confidence; n=85), the median difference in DRAGEN-estimated diploid copy number vs. Bionano was 0.50 copies, and the median difference of our pipeline vs. Bionano was 0.31 copies.

- For allele-specific KIV-2 copy number estimation, we observed very similar performance: RMSE=0.73 copies for DRAGEN (Supplementary Fig. 4C) versus RMSE=0.48 copies from our method (Supplementary Fig. 4D). The improved accuracy of our approach was expected given the enhancements that we implemented: (i) denoising WGS read-depth-derived diploid KIV-2 copy number estimates using genome-wide GC profiles and genomic wave PCs, (ii) partitioning diploid copy numbers into allele-specific estimates using 5 PSVs rather than 2 PSVs, and (iii) handling incomplete intra-allelic homogeneity of PSVs ("exceptions").

We note that these benchmarks of our approach in 1000G indicate modestly higher noise than we estimated in UKB (RMSE=0.4 copies); this was also expected given that in UKB, we applied an enhanced PCA-based denoising approach (Supplementary Note 2) and had a much larger cohort with which to perform phasing and imputation of PSV exception counts. Nonetheless, the strong performance of the approach in 1000G (RMSE=0.48 copies for allele-specific length estimates) indicates its effectiveness even when applied in a simpler, more-easily implemented form, using basic PCA-based read-depth denoising and imputing PSV exception counts from just 546 hom-derived samples spanning the five 1000G superpopulations.

#### 5 Estimating copy-number mutation rate of the KIV-2 repeat

The KIV-2 repeat is understood to be prone to expansion and contraction given its prolific copy-number variation in human populations. However, little is known about its mutational dynamics: how frequently KIV-2 repeat alleles experience copy-number mutations, whether mutations tend to involve small vs. large jumps and expansions vs. contractions, and how the lengths of KIV-2 alleles affect their mutability. One example of a de novo mutation (an intra-allelic loss of 9 repeats from an 18-repeat allele) was observed in an early study that used pulsed-field gel electrophoresis to size KIV-2 alleles representing 376 meioses[7,31]. However, despite remarkable advances in high-throughput genomics technologies and large-scale sequencing of biobank cohorts in the decades since, the mutational dynamics of the KIV-2 repeat have remained very challenging to explore. The main obstacles are the size and intra-allelic homology of the KIV-2 repeat, which preclude accurate measurement of KIV-2 allele lengths using standard short- or long-read sequencing technologies. This problem can be overcome using ultra-high molecular weight sequencing technologies, but such approaches have not been deployed at the scales needed to directly assess KIV-2 mutational dynamics (>10,000 parent-child duos, or ideally >100,000 such duos).

An alternative way to estimate mutation rates is to leverage IBD-sharing among very distantly related individuals within population cohorts, which allows counting mutations not directly observed but rather accrued across many generations of descent from a common ancestor. This approach has been used to estimate the rate of single-nucleotide mutations[102–106], and we recently extended the approach to estimate allele-specific germline mutation rates of short tandem repeats (STRs)[41]. Our analyses of STR mutation rates relied on having high-confidence calls of discreete allele lengths, so the approach was not immediately applicable here: our PSV-based allele length estimates, though accurate, were still noisy (RMSE=0.4 repeat units), such that distinguishing true mutations from measurement noise was still a challenge. However, with two additional innovations (described below), we could adapt the approach to enable a first quantitative exploration of the mutational dynamics of KIV-2 repeat length.

**Estimating overall copy-number mutation rate and jump size distribution.** Our initial examination of allele length concordance between pairs of IBD2 siblings found evidence of a few large mutations (Fig. 2A), but our WGS-derived, PSV-based KIV-2 length estimates were not sufficiently accurate to discern smaller expansions and contractions of 1–2 repeats that were presumably present in the data set. However, visualizing the IBD2 sibling data set provided a key insight: whereas the discordant pairs were informative about large de novo mutations, the strongly concordant pairs—comprising much of the data set—provided an opportunity to identify alleles that could be sized with high confidence based on two concordant length estimates (Fig. 2C). Moreoever, we could augment this set of confidently-sized alleles by expanding this approach beyond IBD2 siblings to include other pairs of related participants.

Concretely, we identified pairs of KIV-2 alleles co-descended from a very recent common ancestor (within the past few generations, such that de novo mutation was very unlikely) from two sources:

1. We searched for such shared alleles among pairs of related individuals (3rd degree or closer) previously identified by UKB[79] based on kinship coefficients computed from SNP-array genotypes[107]. We identified such alleles based on large tracts of shared SNP-haplotypes at *LPA* (specifically, >5 cM of sharing, with >0.5 cM beyond each flank of KIV-2, as computed by hap-ibd[108]). We sorted the identified haplotype pairs in descending order of shared tract length.

2. We augmented this list with a second source of very recently co-descended allele pairs: all pairs of haplotypes with >30 cM of sharing (and >0.5 cM on each flank of KIV-2) with no requirement on kinship; we obtained these pairs from lists of longest shared haplotypes (top "haplotype neighbors") that we previously generated[109]. We also sorted this list in descending order of shared tract length and appended it to the first list.

From among these pairs of very recently co-descended KIV-2 alleles, we identified a subset with concordant PSV-based allele length estimates that converged on a high-confidence, integer allele length call. Specifically, we required that the two continuous-valued PSV-based allele length estimates differ by <1 repeat unit and that their mean be within 0.2 repeat units of an integer. For such allele pairs, we called their length by rounding the mean to this nearby integer. We ensured that no allele length estimate contributed to more than one pair by proceeding iteratively through our list of co-descended allele pairs, excluding a haplotype from further consideration once it had contributed its allele length estimate to an integer copy-number call for a selected pair. This procedure generated a set of ∼10,000 allele pairs with high-confidence copy-number calls.

To estimate the copy-number mutation rate of the KIV-2 repeat, we analyzed IBD-sharing between alleles in this high-confidence KIV-2 copy-number call set (Fig. 2D). Specifically, we arbitrarily selected one SNP-haplotype as a representative from each allele pair in the call set, and we identified IBD tracts between pairs of the selected haplotypes, requiring >3 cM of sharing (and >0.5 cM on each flank of KIV-2) as computed by hap-ibd. This 3-cM cutoff corresponded to a

TMRCA $\sim$88 generations ago (using the derivation in Supplementary Note 3.3, step 4 from ref.[41], with autosomal $N_e(t)$ values that had previously been estimated by analyzing 10,000 White British UKB participants[110]). As such, the IBD data set primarily consisted of pairs of alleles separated by 20–200 meioses (2$\times$TMRCA). Importantly, this separation was large enough that mutation during such a span of generations was not unusual (such that true discordances representing real mutations far outnumbered false positives created by erroneous copy-number calls—see below) but small enough that the probability of observing two mutations within such a span was low.

Comparing the observed rates of copy-number discordance among alleles in an IBD pair (as a function of the number of meioses separating the alleles) to calibration curves indicating expected discordance rates assuming different mutation rates $\mu$, we obtained an estimate of 1 copy-number mutation per $\sim$500 generations (Fig. 2E). In detail, we estimated the number of meioses separating alleles in an IBD pair as $m = 2 \times (\text{TMRCA} - 1)$ (where we subtracted 1 because each high-confidence copy-number call had been generated from a pair of very recently co-descended alleles assumed not to have mutated in the 1+ generation up to their common ancestor), and we estimated the calibration curves as $\Pr(\text{discordance}) = 1 - e^{-m\mu}$. This formula assumes that the likelihood of two or more copy-number mutations that cancel out is negligibly low, which is the case in the parameter space indicated by the data.

We note that the shape of the observed curve (Fig. 2E) provides confidence that the error rate in the high-confidence copy-number call set was well-controlled. The observed fraction of discordant pairs increased roughly linearly with $m$, as expected, whereas if a large fraction of the discordances arose from errors in copy-number calls, this would inflate the discordance rates by a constant offset irrespective of the IBD tract length (and hence the number of meioses) between two alleles. We also note that the pairwise discordances analyzed here are likely to represent a largely nonredundant set of mutational events given that the 369 discordant pairs that were observed involved 478 unique alleles (haplotypes).

We estimated the jump-size distribution of KIV-2 copy-number mutations using the empirical distribution of observed pairwise discordances (Fig. 2F). This is inexact because a small fraction of the pairwise discordances are expected to represent two or more mutations, but the fraction is small enough not to affect the qualitative conclusions, which were clear: most KIV-2 copy-number mutations are 1-unit expansions or contractions, 2–4-unit jumps are also fairly common, and larger jumps are rare but occur as well.

**Estimating allele-specific large-jump expansion and contraction rates.** The above approach provided an overall estimate of the frequency of de novo copy-number mutations and a glimpse at the jump-size distribution of such mutations, but the analytical approach could not distinguish expansions from contractions (because the source allele of discordant pairs was unknown), and the set of mutations visible among the high-confidence copy-number calls was far too small to explore the relationship between allele length and mutability. To overcome these limitations, we devised a second, complementary approach that focused on exploring the mutational dynamics of large-jump mutations (i.e., multi-copy expansions and contractions).

The key intuition of this approach was that if we focused specifically on analyzing large-jump mutations, we no longer needed exact copy-number calls: the approximate (RMSE=0.4 repeat units) PSV-based allele length estimates that we had generated for nearly half of the UKB cohort (215,936 individuals) were sufficiently accurate for this purpose. This much larger haplotype set was suitable for analysis using the approach we previously developed for estimating allele-specific germline mutation rates of STRs[41]. In brief, the approach works by identifying pairs of alleles forming two-leaf clades on the coalescent tree (identified based on being mutually-top haplotype neighbors[41,109]) and imputing their ancestral allele from other haplotypes in the cohort (Fig. 2G). Comparing the extant two alleles (from the IBD pair) to their imputed ancestral allele then provides an opportunity to observe mutations along the the lineage connecting the IBD pair, and any such mutations can be determined to be expansions or contractions based on the length of the imputed ancestral allele.

To implement this approach, we first identified mutually-longest IBD matches using the lists of top haplotype neighbors that we had previously generated[109], restricting to individuals with PSV-based allele length estimates and requiring IBD tracts to have length >3 cM with >0.5 cM on each flank of KIV-2. Among this set of IBD pairs, we identified those with large discordances between their estimated allele lengths (≥3 repeat units). We imputed ancestral alleles of such pairs from the remaining PSV-phased haplotypes (taking the average of the two allele lengths imputed into the two members of the pair, which were usually very similar). To maximize the accuracy of these imputed ancestral alleles, we further restricted the analysis to EUR-ancestry individuals (n=203,720 with PSV-based allele length estimates).

Among the mutually-longest IBD pairs in this data set with highly discordant allele lengths (differing by ≥3 repeat units), suggesting a large-jump mutation, we filtered to those for which the observed and ancestral (imputed) allele lengths were consistent with a single mutation on one branch of the two-leaf clade. Specifically, we required that one extant allele's PSV-based length estimate differ by >2 repeat units from the imputed ancestral allele (strongly suggesting a mutation) and that the other allele be <1.5 repeat units from the imputed ancestral allele (consistent with no mutation). These criteria defined a subset of the mutually-longest IBD pairs with a high-confidence multi-copy mutation of known directionality (expansion or contraction).

To estimate allele-specific large-jump mutation rates (Fig. 2H), we computed the total number of expansions (respectively, contractions) that the above procedure identified among mutually-longest IBD pairs with ancestral alleles of a given length (after rounding to the nearest integer), and we divided this number by the total length of IBD lineages across all mutually-longest IBD pairs with ancestral alleles of that length (obtained for non-discordant pairs by rounding the average of the pair's SNP-phased allele length estimates to the nearest integer). The rationale for this computation was that each such ancestral allele had had the opportunity to experience a large-jump mutation in any of 2×TMRCA meioses that could have created a detectable allele-length discordance. We estimated TMRCA in a manner similar to our previous work[41], except that here we adjusted upfront for ascertainment of longest IBD matches (rather than implementing a correction downstream[41]). To do this, we used a similar coalescent approach to estimate the expected time to first coalescence with any of a set of $n$ lineages (rather than the time to first coalescence with a single lineage, which we previously computed[41]). This required modifying the computation of $P(\text{TMRCA} = t)$ as follows. We initialized $n$ to $2 \times 203,720 - 1$ (the number of other PSV-phased haplotypes in the analysis), and at each generation $t$, we updated $n \leftarrow 2N_e(t) \cdot \left(1 - (1 - \frac{1}{2N_e(t)})^n\right)$ (because each of the $2N_e(t)$ haplotypes in the previous generation had a $(1 - \frac{1}{2N_e(t)})^n$ probability of not being an ancestor of any current haplotype). We then computed the probability of coalescence with one of the haplotypes in the previous generation as $p_{\text{coal}} = n/(2N_e(t))$.

Some caveats we note are that the actual lengths of the putative large-jump mutations considered in this analysis are uncertain: we ascertained IBD pairs with discordances of ≥3 repeat units between their continuous-valued estimated lengths, but owing to noise in the PSV-based length estimates, some of the mutations could have been shorter, and a small fraction could have been false positives. False negatives were also possible: e.g., a genuine 3-repeat-unit mutation would be equally likely to create an observed discordance of greater or less than 3 repeat units. As such, the absolute estimates of large-jump mutation rates (and the meaning of "large-jump" in this context) are complicated to interpret. Nonetheless, the qualitative conclusions from the results of the analysis should be robust to these issues and suggest that false-positive mutation calls were well-controlled: random errors arising from erroneous detections would be expected to be balanced between expansions and contractions, unlike the pronounced contraction bias observed for longer alleles (Fig. 2H).

### 6 Factors limiting SNP-based prediction of Lp(a)

In general, polygenic score transferability across ancestries is limited by differences in linkage disequilibrium (LD) patterns between causal variants and variants used for prediction[111]. Strategies that prioritize likely-causal variants (e.g., variants supported by statistical fine-mapping) tend to exhibit higher accuracy across populations[112]. To better understand the particularly low transferability of SNP-based Lp(a) predictions, we examined the haplotypes of two common SNPs (rs10455872 and rs3798220) that were among the first to be associated with high Lp(a) (ref.[2]). These two SNPs have frequently been used to predict high Lp(a) and to stratify cardiovascular risk[2,113,114]; however, whereas rs10455872>G associates with high Lp(a) across genetic ancestries, rs3798220>C associates with high Lp(a) only in EUR-ancestry cohorts[50].

Examination of KIV-2 alleles on phased UKB haplotypes carrying these variants explained this behavior (Fig. 3B): for each SNP, most EUR-ancestry haplotypes carrying the risk allele also carried short (10–12 repeat) KIV-2 alleles (which encode efficiently-secreted apo(a) isoforms and increase Lp(a) levels[15]), such that each of the two SNPs associated with high Lp(a) (130.5 and 155.5 nmol/L) among individuals of European genetic ancestry. For rs10455872, this pattern extended to other ancestries, but for rs3798220, non-EUR haplotypes carrying the risk allele tended to contain much longer KIV-2 alleles (median 21–25 repeat units) and associate with much lower Lp(a) (mean 9.5–57.1 nmol/L). The weak LD between rs3798220 and KIV-2 length in non-EUR ancestries is primarily explained by their genomic distance (71 kb; 0.058 cM): recombination randomizes the KIV-2 allele present on a rs3798220 haplotype every $\sim$1,700 generations (a time scale shorter than the divergence time of EUR and AFR haplotypes[115]. In contrast, rs10455872 is much closer to the KIV-2 repeat region (0.006 cM), such that it tags short KIV-2 alleles much more stably. The rapid mutation of the KIV-2 repeat (once per $\sim$500 generations; Fig. 2E) further contributes to breakdown of LD between SNPs and KIV-2 (independent of genetic distance).

In addition to the difficulty of tagging the highly multiallelic, highly mutable KIV-2 repeat with SNPs, another factor limiting SNP-based prediction of Lp(a)—and potentially other phenotypes—is the presence of strong *cis*-epistatic interactions among causal variants on a haplotype. We previously identified several common variants across *LPA* that are likely to causally influence a haplotype's Lp(a) production, and these variants span the entirety of *LPA* from its 5' UTR (rs1853021 and rs1800769) to its final exon (rs3124784; *LPA* R2016C)[26]. Because these variants have approximately multiplicative effects on the amount of Lp(a) produced by a haplotype[26] (Fig. 3C), cross-ancestry prediction of Lp(a) using linear, SNP-based models requires model variants to tag entire *LPA* haplotypes (which span 132 kb and recombine every $\sim$1,500 generations) in all populations. This is not possible as common, globally-polymorphic SNPs are typically many thousands of generations old[116]. This limitation extends to linear polygenic prediction of any trait for which at key loci, multiple causal variants on a haplotype exert *cis*-epistatic effects on phenotype.

### 7 Estimating Lp(a) heritability attributable to the *LPA* locus

To estimate the maximum accuracy achievable by an optimal *LPA*-based genetic predictor of Lp(a) (grey shaded rectangles in Fig. 3A), we estimated the fraction of Lp(a) variance attributable to genetic variation at the *LPA* locus using sibling data in UKB. Our approach is similar to Falconer's method for estimating heritability in twin studies[117]. We regard Lp(a) as a sum of three independent random variables:

$$\text{Lp(a)} = G_{LPA} + C + E. \tag{17}$$

Here, $G_{LPA}$ is the genetic component of Lp(a) due to variation at *LPA*, $C$ is the remaining component shared by siblings, and $E$ captures the remaining influences on Lp(a) not shared among siblings. Each of the components $C$ and $E$ include non-genetic as well as genetic influences due to variation at loci other than *LPA*. We write $\sigma^2_{LPA}$, $\sigma^2_C$, and $\sigma^2_E$ for the respective variances of these components, and we are interested in the portion of total Lp(a) variance ($\sigma^2_{\text{Lp(a)}}$) attributable to *LPA*:

$$H^2_{LPA} := \sigma^2_{LPA}/\sigma^2_{\text{Lp(a)}}. \tag{18}$$

We can express the correlation among pairs of siblings who share $k$ alleles IBD at *LPA* as:

$$\text{IBD}_k R_{\text{Lp(a)}} = \left(\frac{k}{2}\cdot\sigma^2_{LPA} + \sigma^2_C\right)/\sigma^2_{\text{Lp(a)}} \tag{19}$$

where $k$ = 0 or 2. (This formula also holds for $k$ = 1 under the assumption that $G_{LPA}$ is additive across the two parental haplotypes at *LPA*.) The quantity $H^2_{LPA}$ can thus be expressed as:

$$H^2_{LPA} = \text{IBD}_2 R_{\text{Lp(a)}} - \text{IBD}_0 R_{\text{Lp(a)}}. \tag{20}$$

To estimate $H^2_{LPA}$ for individuals of European genetic ancestry, we applied equation (20) directly to EUR siblings in UKB. We identified 3,171 non-overlapping EUR-ancestry IBD2 sibling pairs with Lp(a) measurements by counting discordant SNP array genotypes in a 4-Mb region surrounding *LPA* (chr6:158.6–162.6 Mb), requiring IBD2 pairs to have $\leq$10 discordances, and we observed:

$$\text{IBD}_2 R_{\text{Lp(a)}} = 0.909,\ 95\%\ \text{CI} = [0.903, 0.915] \quad \text{(between EUR siblings)}.$$

Since we evaluated Lp(a) prediction accuracy among individuals with PSV-based KIV-2 phasing estimates, we also computed sibling correlation for IBD2 siblings with PSV-based KIV-2 phasing estimates and observed a similar correlation:

$$\text{IBD}_2 R_{\text{Lp(a)}} = 0.900,\ 95\%\ \text{CI} = [0.890, 0.910] \quad \text{(between PSV-phased EUR siblings)}.$$

We next identified 3,905 non-overlapping EUR-ancestry IBD0 sib pairs by counting discordant homozygous WGS genotypes in the region chr6:160.42–160.78 Mb (excluding variants in the KIV-2 region, variants with MAF<0.01, and variants with AAScore<0.8), requiring IBD0 sibs to have $\geq$2 discordances. Correlation of Lp(a) estimates between IBD0 siblings was low with a wide confidence interval (95% CI = [–0.007, 0.056]), so we leveraged accurate genetic predictions of Lp(a) (derived from SNP-based KIV-2 phasing) to obtain a more precise estimate:

$$\text{IBD}_0 R_{\text{Lp(a)}} = \text{IBD}_0 R_{\text{resid Lp(a)}} \cdot \sigma^2_{\text{resid Lp(a)}}/\sigma^2_{\text{Lp(a)}} = 0.012, \quad 95\%\ \text{CI} = [0.008, 0.016],$$

where resid Lp(a) = measured Lp(a) – predicted Lp(a).

Combining these estimates for $\text{IBD}_2 R_{\text{Lp(a)}}$ and $\text{IBD}_0 R_{\text{Lp(a)}}$, we obtained:

$$H^2_{LPA} = 0.888 \quad \text{(among PSV-phased EUR individuals)}$$
$$H^2_{LPA} = 0.897 \quad \text{(among all EUR individuals).}$$

We needed a different approach to estimate $H^2_{LPA}$ in non-EUR ancestries: for these ancestries, the UKB cohort contained relatively small numbers of sibling pairs (326 SAS, 222 AFR, and 26 EAS sibling pairs), such that direct application of equation (20) yielded estimates of $H^2_{LPA}$ with confidence intervals too wide to obtain a useful upper bound on prediction accuracy. Instead, we estimated $\text{IBD}_2 R_{\text{Lp(a)}}$ in non-European ancestries using data from EUR siblings, subsampling to match each ancestry's Lp(a) distribution (so as to account for the effect of heteroskedasticity in Lp(a) measurement noise on $H^2_{LPA}$). In each analyzed non-EUR ancestry group, we stratified individuals with PSV-based phased KIV-2 length estimates into 50 equal-sized bins (each representing 2% of the Lp(a) distribution in that ancestry). For each bin, we chose 20 EUR PSV-phased IBD2 sibling pairs for which the first sibling's measured Lp(a) (using a random ordering of the two siblings) was within the bin. We estimated each ancestry's $\text{IBD}_2 R_{\text{Lp(a)}}$ as the median observed sibling correlation across 100 runs of this experiment, and we estimated $H^2_{LPA}$ using equation (20), assuming an equal portion of variance was explained by $C$ across ancestries (i.e., $\text{IBD}_0 R_{\text{Lp(a)}} = 0.012$ in each ancestry). In this way, we obtained the following estimates for $H^2_{LPA}$ used in Fig. 3A:

$$H^2_{LPA} = 0.888 \text{ (EUR)}, 0.827 \text{ (AFR)}, 0.876 \text{ (EAS), and } 0.866 \text{ (SAS).}$$

We note that our model of Lp(a) in equation (17) excludes interactions between the variance components, which likely contribute to some degree to Lp(a) measurements (e.g., as evidenced by heteroskedasticity in Lp(a)). The variance contributed by interactions between $G_{LPA}$ and $C$ will be reflected in $\text{IBD}_2 R$ but not $\text{IBD}_0 R$, resulting in a slight overestimate of $H^2_{LPA}$ derived from equation (17). We expect interaction-related inflation is modest given that our model for Lp(a) derived from *LPA* genetic variation explains >95% of estimated $H^2_{LPA}$ in individuals of European genetic ancestry.

#### 8 Genetic architecture of Lp(a) across ancestries

The key contributors to the genetic architecture of Lp(a) across ancestries represented in UKB can be summarized as follows:

- In all ancestries, the KIV-2 length polymorphism plays a primary role in shaping Lp(a).
- In EUR and SAS, two ancestries with similar KIV-2 length distributions, Lp(a) is further shaped by seven common, large-effect protein-altering variants that reduce Lp(a) by >90%. Collectively, these variants suppress a large fraction (48% in EUR, 47% in SAS) of short and medium-length KIV-2 alleles (≤15 repeat units). Three smaller-effect common SNPs play a secondary but important role in shaping Lp(a).

- In AFR, an ancestry with a broadly similar KIV-2 length distribution, the large-effect Lp(a)-reducing variants common in EUR have much lower frequencies (combined AFR AF=3%), and Lp(a) is further shaped by a common cryptic splice variant (rs41270998>G, AFR AF=7% vs. EUR AF=0.2%) and by the high frequency of a common, smaller-effect Lp(a)-increasing promoter variant (rs1800769>T, AFR AF=47% vs. EUR AF=17%; 25% increase in Lp(a)).

- In EAS, Lp(a) is shaped by a relative dearth of short and medium-length KIV-2 alleles ($\leq$15 repeat units; combined EAS AF=9% vs. EUR AF=29%), of which 9% segregate with the common, EAS-specific canonical splice variant in KIV2 (EAS AF=1.1% vs. <0.06% in other populations) and produce little or no Lp(a).

### 9 Lp(a) prediction and cardiovascular diseases in AoU

We generated genetic predictions of Lp(a) in AoU (v7) to assess the feasibility of using predicted Lp(a) to stratify cardiovascular disease risk out-of-sample and across diverse populations.

AoU is a longitudinal cohort of participants aged 18–90+ years at recruitment[118]. Most participants provided biosamples and data from surveys and electronic health records (EHR); data on subsequent health outcomes were obtained from EHR linkage. Short-read WGS data were generated from blood-derived DNA for 245,394 individuals in the AoU v7 release. Sequencing was performed using Illumina NovaSeq 6000 machines and reads were aligned to GRCh38 with Illumina DRAGEN Bio-IT Platform Germline Pipeline 3.4.12 as previously described[34]. Samples were sequenced to genome-wide average coverage $\geq 30\times$.

We broadly followed the strategy we employed in UKB for generating *LPA* KIV-2 + SNP haplotypes and Lp(a) predictions, with a few differences detailed below.

1. **Correcting for GC bias through analysis of allelic depths in VCF files.** We estimated each WGS sample's GC bias profile by analyzing allelic depth data provided in population level VCF files available for the AoU cohort (acaf_threshold_v7.1). This approach avoided analysis of read-level CRAM files for GC profile estimation, which was feasible for UKB but much more expensive for AoU owing to charges for accessing data from nearline storage. For each sample, we computed site-level read depths as sums of allelic depths (across alleles of a given variant), which were reported in the "AD" field for each variant for which an individual was heterozygous or homozygous for an alternate allele. We masked sites in low-complexity regions, regions prone to mapping errors, and commonly copy-number-variable regions as in our analysis in UKB. To avoid double-counting reads derived from the same DNA fragment, we restricted to variant sites (of a given individual) that were separated by $\geq$400 bp. We also restricted to biallelic sites with allele frequency >0.05 and excluded indels.

For each remaining SNP site, we computed its GC context as the GC content of the 400-bp region of GRCh38 centered at the site. We then computed the sample's GC-response curve in units of read-depth-per-bp (for each possible value of GC context) as:

$$B(gc) = \frac{\sum_{\text{sites with GC context}=gc} (\text{read depth})}{\#\ \text{sites with GC context} = gc}, \quad 0 \leq gc \leq 400.$$

We next estimated the KIV-2 allele length sum (dipCN) by extracting read alignments to *LPA* from the CRAM file, counting read alignments with starting position within chr6:160611533–160646720, weighting each read by the inverse of the GC-response curve, dividing by the length of the KIV-2 repeat unit (5,544bp), and normalizing against the analogous quantity computed in the non-copy-number-variable regions flanking KIV-2 (chr6:160.2–160.6 Mb and chr6:160.65–161.05 Mb, excluding masked sites):

$$\text{dipCN} = 2 \cdot \frac{\sum_{gc=0}^{400} \sum_{\text{KIV-2 reads with GC context}=gc} \frac{1}{B(gc)}}{\text{KIV-2 repeat unit length}} \cdot \frac{\text{length of unmasked flank region}}{\sum_{gc=0}^{400} \sum_{\text{flank reads with GC context}=gc} \frac{1}{B(gc)}}.$$

For the GC context of a read, we used the GC content of the 400-bp window in GRCh38 with leftmost endpoint starting at the alignment start position. We adjusted these dipCN estimates by a constant additive offset (–2×0.36 repeat units, in agreement with visual inspection of the peaks of the dipCN histogram) to adjust for the overlap of the analyzed region (chr6:160611533–160646720) with KIV-1 and KIV-3.

2. **Genotyping PSV copy number.** We genotyped PSVs in and near exons of KIV-2 using samtools mpileup following the procedure described for UKB (Methods).

3. **Generating an *LPA* SNP haplotype scaffold.** We extracted SNP-array genotypes of common variants with MAF>0.01 and missingness <0.1 in the ∼20 Mb region surrounding KIV-2 (chr6:150601532–170656869) using plink2. We phased the SNP-array genotypes using SHAPEIT5 v5.1.1 (phase_common) to form a haplotype scaffold.

4. **Phasing KIV-2 dipCN and common PSVs.** We phased KIV-2 dipCN and common PSVs using our SNP-based phasing algorithm (leveraging shared haplotypes within AoU).

5. **Phasing rare PSVs and rare Lp(a)-modifying SNPs.** We phased the remaining biallelic variants used in our model onto the scaffold of SNP-array haplotypes using SHAPEIT5 v5.1.1 (phase_rare).

6. **Predicting Lp(a) from *LPA* diplotypes.** We predicted each individual's Lp(a) using the resulting phased haplotypes and the model parameters estimated from UKB (Supplementary Tables 2 and 3).

We tested genetically predicted Lp(a) for association with chronic ischemic heart disease (ICD-10 code I25). We restricted analysis to unrelated AoU participants, randomly dropping one individual from each related pair with kinship coefficient >0.0884, and separately analyzing the three most prevalent genetic ancestries (`eur`, `afr`, and `amr`). Genetic ancestry assignments and kinship coefficients were obtained from the AoU v7 Controlled Tier Curated Data Repository (CDR). We identified I25 cases using the AoU Cohort Builder to identify individuals whose health record contained a condition code categorized as chronic ischaemic heart disease (SNOMED ID 315286). We used logistic regression to test predicted Lp(a) for an association with I25, including as covariates age (as of 01/01/24), age$^2$, and self-reported sex at birth. To determine odds ratios for Lp(a) ranges relative to low Lp(a) (<30 nmol/L), we binned individuals according to their predicted Lp(a) (bin cutoffs 0, 30, 100, 200, and 600 nmol/L). We performed logistic regression analysis including indicator variables for membership in bins [30, 100), [100, 200), and [200, 600) with covariates age, age$^2$, and sex, interpreting parameter estimates for coefficients of the indicator variables as log odds ratios.

#### 10 Genome-wide association analysis of Lp(a) adjusted for *LPA* variation

Searching for genome-wide genetic influences on Lp(a) has been complicated by the large amount of Lp(a) variance contributed by genetic variation at *LPA*[119]. Here, our model of Lp(a) together with accurate genotyping and phasing of sequence variation at *LPA* (including the KIV-2 repeat) in UKB allowed us to almost completely control for effects of *LPA* variation on Lp(a), greatly increasing statistical power to detect genetic effects on Lp(a) at other loci.

In detail, we computed an *LPA*-adjusted Lp(a) phenotype by taking the ratio of observed-to-predicted Lp(a) levels (cropping each from below at 3.8 nmol/L) and applying rank-based inverse normal transformation:

$$\frac{\max(3.8, \text{measured Lp(a)})}{\max(3.8, \text{predicted Lp(a)})} \stackrel{\text{INT}}{\longmapsto} \textit{LPA}\text{-adjusted Lp(a)}. \qquad (21)$$

In this formula, measured Lp(a) and predicted Lp(a) are on the "raw" scale with a mean of 1.51 nmol/L in null-allele homozygotes as in equation (6).

We used BOLT-LMM to perform GWAS on common and low-frequency variants imputed in UKB EUR-ancestry participants, including sex, age, age squared, and 20 genetic principal components as covariates (n=424,463 participants with no missing data and who had not withdrawn from the study; we did not exclude related individuals, as linear mixed model association analysis is robust to sample relatedness[120]). We ran BOLT-LMM on the imp_v3 variant set previously generated by UKB using the HRC and UK10K+1KGP3 imputation reference panels[79]. We restricted analysis to 17.1 million variants with MAF>0.001 and imputation INFO>0.3.

To define loci and identify index variants associated with *LPA*-adjusted Lp(a) ($P<5\times10^{-8}$), we computed linkage disequilibrium between all pairs of nearby significantly-associated variants using plink2 alpha 5.22 --r-unphased ref-based --ld-window-kb 3000 --ld-window-r2 0.0001. We then identified a subset of pairwise-independent associations by iterating through variants in descending order of association strength and retaining a variant only if it remained significantly associated with *LPA*-adjusted Lp(a) ($P<5\times10^{-8}$) in conditional analysis with each previously-retained nearby variant[60]. Finally, we defined loci and index variants by iterating through the remaining variants (again in descending order of association strength) and retaining a variant only if it was >500 kb away from any previously-retained variant and >1Mb away from any previously-retained very strongly associated variant ($P<10^{-100}$). This procedure identified 330 index variants independently associated ($P<5\times10^{-8}$) with *LPA*-adjusted Lp(a) and separated by at least 500 kb. The expected number of false positives at a significance threshold of $P<5\times10^{-8}$ is <1, such that this conventional genome-wide significance threshold provided adequate control of false discoveries.

We annotated index variants with protein-coding genes within 100 kb based on GENCODE[121] v35 (lifted to GRCh37). We annotated coding consequences of variants (and of significantly-associated variants in strong LD with index variants; $R^2>0.9$) as previously described[60]. We annotated index variants reported to associate with gene expression (eQTLs)[122] using eQTL data downloaded from the GTEx Portal on 08/17/2021.

To explore genetic overlap between *LPA*-adjusted Lp(a) and other lipid measurements, we collated summary association statistics for *LPA*-adjusted Lp(a) with summary statistics for LDL, HDL, total cholesterol, and triglyceride levels from our previous analyses in UKB[95]). To generate scatter plots comparing effect sizes for other lipid traits to effect sizes for *LPA*-adjusted Lp(a), we identified independent index variants for each lipid phenotype using the procedure described above. For a given lipid phenotype, we then merged its index variants with the set of index variants for *LPA*-adjusted Lp(a) and LD-pruned the merged list (removing the variant with the weaker association among pairs of variants with $R^2>0.5$). We computed SNP-heritability and genetic correlations between *LPA*-adjusted Lp(a) and other lipid traits using BOLT-REML[123], using the same covariates and sample exclusions as for BOLT-LMM and additionally downsampling the data 8-fold and using the --remlNoRefine option for computational efficiency (as additional accuracy was not needed).

We identified 92 index variants with effect sizes for *LPA*-adjusted Lp(a) greater than twice as large in magnitude (in *z*-score units) as their effect sizes on any of the four other lipid phenotypes. We performed gene ontology (GO) enrichment analysis (https://geneontology.org/)[124–126] on the set of genes nearest to these index variants (including multiple genes if an index variant overlapped multiple genes). GO enrichment analysis identified enrichment for a single Reactome pathway, "Synthesis of PC" (phosphatidylcholine) that passed multiple hypothesis correction ($P=1.43\times10^{-5}$, Bonferroni-adjusted $P=0.034$).

##### Exome-wide association analysis of rare coding variant effects on Lp(a)

We additionally performed exome-wide association analysis of *LPA*-adjusted Lp(a) by running BOLT-LMM on variants called from whole-exome sequencing (WES) of UKB[127] and on WES-derived pLoF burden "variants" (n=409,572 participants after restricting to individuals with WES data). Specifically, we extracted pLoF annotations from gnomAD v4.1[128] and identified variants with a high-confidence LoF annotation for the MANE Select transcript[129] of a protein-coding gene, no LoF_filter or LoF_flags, FILTER=PASS, and >90% call rate in the UKB WES data set. We then combined these pLoF variants with pLoF CNV calls[95] into burden masks using regenie[130] v3.1.1, restricting to variants with alternate AF<0.001. We restricted pLoF burden analyses to 15,827 genes with a combined pLoF allele count of at least 5 among individuals included in analysis. The main manuscript describes associations that reached a conservative significance threshold of $P<10^{-6}$, and in Supplementary Table 5 we report additional associations that reached FDR<0.05.

We restricted single-variant association analyses of WES variant calls to 3.2 million variants with AF>$2\times10^{-5}$, and we excluded variants within 5 Mb of *LPA*. Among Bonferroni-significant associations ($P<1.6\times10^{-8}$), we identified variants with large effect sizes (0.5 standard deviations or larger), reported in Supplementary Table 6. Fig. 4E,F and Supplementary Fig. 6 show data for the subset of these variants that either affected genes not in the pLoF burden $P<10^{-6}$ set or had effect directions opposite to pLoF effects.

To compute corresponding effect sizes and association strengths for other lipid phenotypes, we ran BOLT-LMM on each of the other lipid phenotypes with the same parameter settings.

#### 11 Testing rare coding, splice region, and promoter variants for association with Lp(a)

To test rare variants in and near exons of *LPA* for association with Lp(a) (after accounting for Lp(a) predictions from our model), we developed a likelihood ratio test capable of modeling the heteroskedasticity of Lp(a) measurements and estimating multiplicative effects of Lp(a)-modifying SNPs. To test a particular SNP (or indel) $s$, we computed the test statistic:

$$\lambda_{LR} = 2 \cdot \log \frac{\sup_{\beta_s} L\left(\text{data} \mid \beta_s\right)}{L\left(\text{data} \mid \beta_s = 1\right)}, \tag{22}$$

where $L$ is the likelihood of the observed data (i.e., measured Lp(a) values and *LPA* haplotypes) given the model parameter ($\beta_s$, the multiplicative effect of SNP $s$). The quantity $\lambda_{LR}$ is asymptotically $\chi^2_1$-distributed under the null hypothesis $\beta_s = 1$.

To apply the likelihood ratio test, we used the observed data to estimate the probability density function for observing a particular Lp(a) measurement given an Lp(a) prediction:

$$P\left(\text{measured Lp(a)} \mid \text{predicted Lp(a)}\right) \tag{23}$$

where measured and predicted Lp(a) are on the "raw" scale as before (i.e., with a mean of 1.51 nmol/L offset in individuals homozygous for null alleles). We worked with cropped and log-transformed raw Lp(a) values, applying the following transformation mapping Lp(a) values to the interval [0, 52]:

$$T\left(\text{Lp(a)}\right) = \begin{cases} 0 & \text{if Lp(a)} < 2 \\ 1 + 50 \cdot \frac{\log_{10}(\text{Lp(a)}/2)}{\log_{10}(400/2)} & \text{if } 2 \leq \text{Lp(a)} < 400 \\ 52 & \text{if Lp(a)} \geq 400. \end{cases} \tag{24}$$

We used the observed data (Lp(a) measurements and Lp(a) predictions from our model, for the set of unrelated EUR-ancestry UKB participants) to first estimate a discretized joint probability distribution of measured and predicted Lp(a). We did so by assigning a mass $M(x, y)$ to each integer grid point in the square [0, 52] x [0, 52], distributing a unit mass associated to each transformed observed data point

$$(x, y) = (T(\text{measured Lp(a)}), T(\text{predicted Lp(a)})) \tag{25}$$

across the four nearest grid points. The portion of mass accruing to the proximal grid point $(x_0, y_0)$ was determined by the formula $(1 - |x - x_0|) \cdot (1 - |y - y_0|)$ .

To estimate the conditional probability distribution of $x = T(\text{measured Lp(a)})$ given $y = T(\text{predicted Lp(a)})$, we normalized the total mass on each horizontal grid line (corresponding to a fixed Lp(a) prediction) to be 1 and regularized the result:

$$P(x \mid y) = 0.95 \cdot M(x, y) / \sum_{i=0}^{52} M(i, y) + 0.05/53. \tag{26}$$

Finally, to extend this discretized conditional probability distribution to non-integer values of transformed Lp(a), we used a bivariate spline (scipy.interpolate.RectBivariateSpline[131] with `kx`=1, `ky`=1) to interpolate between grid points. The resulting spline fit provided an approximation of the conditional probability density $P\left(\text{measured Lp(a)} \mid \text{predicted Lp(a)}\right)$.

We tested all SNP and indel variants in or near an *LPA* exon or within the *LPA* promoter, using the empirically estimated conditional distribution $P\left(\text{measured Lp(a)} \mid \text{predicted Lp(a)}\right)$ to compute the likelihood ratio test statistic $\lambda_{LR}$ (comparing the likelihood of the observed data under a model in which the variant was allowed to influence predicted Lp(a) vs. the likelihood under the model in which the variant had no effect). We tested all rare variants (EUR AF<0.001) within 20 bp of an *LPA* exon or in the 1-kb region upstream of the transcription start site (to conservatively cover the *LPA* promoter). For each such SNP or indel $s$, we restricted analysis to individuals with a haplotype carrying $s$ that was predicted under our model (assuming no effect of $s$) to contribute $\geq$5 nmol/L of Lp(a), and we required $\geq$5 such carriers among unrelated EUR participants. To allow all rare variants to be considered equally within this testing framework, we computed Lp(a) predictions using a slightly modified model in which we excluded variants with EUR AF<0.001 that we had included in the model we used for other analyses (Supplementary Table 3), such that these variants were allowed to be rediscovered by this test.

For each SNP or indel $s$ meeting these criteria, we evaluated the likelihood of the observed data assuming different possible values of the multiplicative effect size $\beta_s$ of the variant. Specifically, for each of a finely-discretized set of values of $\beta_s$, we computed Lp(a) predictions among carriers of $s$ (augmenting our model with the multiplicative effect $\beta_s$ of variant $s$) and computed the likelihood

$$L(\text{data} \mid \beta_s) = \prod_{\text{carriers of s}} P\left(\text{measured Lp(a)} \mid \text{predicted Lp(a) with SNP } s \text{ effect} = \beta_s\right) \tag{27}$$

of the observed Lp(a) measurements under this model. We evaluated values of $\beta_s$ ranging from –5 to 5 (using a step size of 0.01), allowing negative values of $\beta_s$ to avoid statistical subtleties that arise when applying a likelihood ratio test to a one-sided parameter space. We determined the maximum likelihood estimate for the effect size $\widehat{\beta_s} = \text{argmax}_{\beta_s} L\left(\text{data} \mid \beta_s\right)$, from which we computed the test statistic

$$\lambda_{LR} = 2 \cdot \log \frac{L\left(\text{data} \mid \beta_s = \widehat{\beta_s}\right)}{L\left(\text{data} \mid \beta_s = 1\right)} \tag{28}$$

and the upper and lower bounds of the $\exp(1.96^2/2)$-likelihood region approximating a 95% confidence interval (used to determine error bars in Fig. 5C). This likelihood region is the set of parameters $\beta$ satisfying:

$$2 \cdot \log \frac{L\left(\text{data} \mid \beta_s = \widehat{\beta_s}\right)}{L\left(\text{data} \mid \beta_s = \beta\right)} < 1.96^2. \tag{29}$$

Finally, we applied a series of conservative filters to remove significant associations that could potentially reflect linkage disequilibrium with other Lp(a)-modifying variants rather than causal biological effects. First, we removed variants with smaller estimated effect sizes ($|\widehat{\beta_s} - 1| < 0.5$), reasoning that some of these associations might reflect LD with common variants not included in our model. Second, we filtered any variant whose association could be largely explained by inclusion of another rare variant in the Lp(a) model. Specifically, for each significantly associated variant $s$ with a large effect size ($P<10^{-5}$ and $|\widehat{\beta_s} - 1| > 0.5$), for each potential tag SNP $t$ with

$\lambda_{LR} > 10$, we recomputed the likelihood ratio test statistic for $s$ after including $t$ in the model for predicting Lp(a), assigning $t$ its maximum likelihood effect size $\widehat{\beta}_t$. We filtered significantly associated SNPs whose association strengths either decreased by $>90\%$ on the $\lambda_{LR}$ scale or were not significantly associated ($P>10^{-5}$) in a joint analysis with another variant $t$. This filtering strategy prioritizes obtaining a high-confidence set of likely-causal variants rather than maximizing the size of the set (e.g., any variant in perfect LD with another variant is removed); several filtered variants were annotated as pLoF or were missense variants with a high PrimateAI-3D score and probably affect apo(a) function despite being filtered (Supplementary Table 7 and Supplementary Fig. 8).

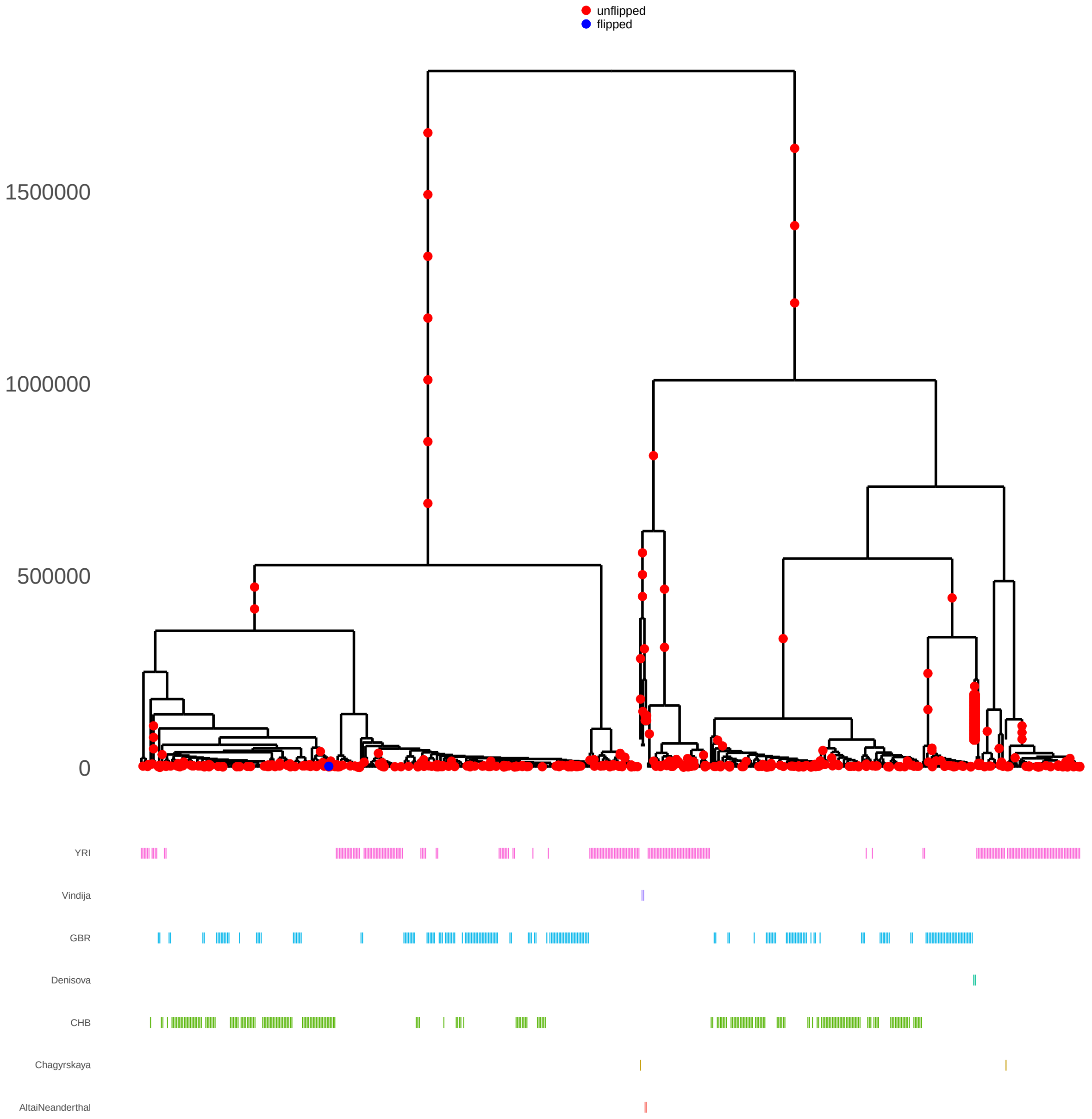


**Supplementary Figure 1. Coalescent tree of modern human and archaic hominin haplotypes at the KIV-2 locus.** This tree was previously inferred by Relate using data from 302 modern human individuals (1000 Genomes CHB, GBR, YRI) and four archaic hominin individuals[35]. Fig. 1A shows a subtree in which the modern human haplotypes were downsampled by a factor of 10. The plots were generated using the Relate TreeView.sh script on the 1000G_sub_Nea data set for the GRCh37 position 6:161071212; the $y$-axis indicates coalescence times in years, assuming 28 years per generation as suggested in the Relate documentation.

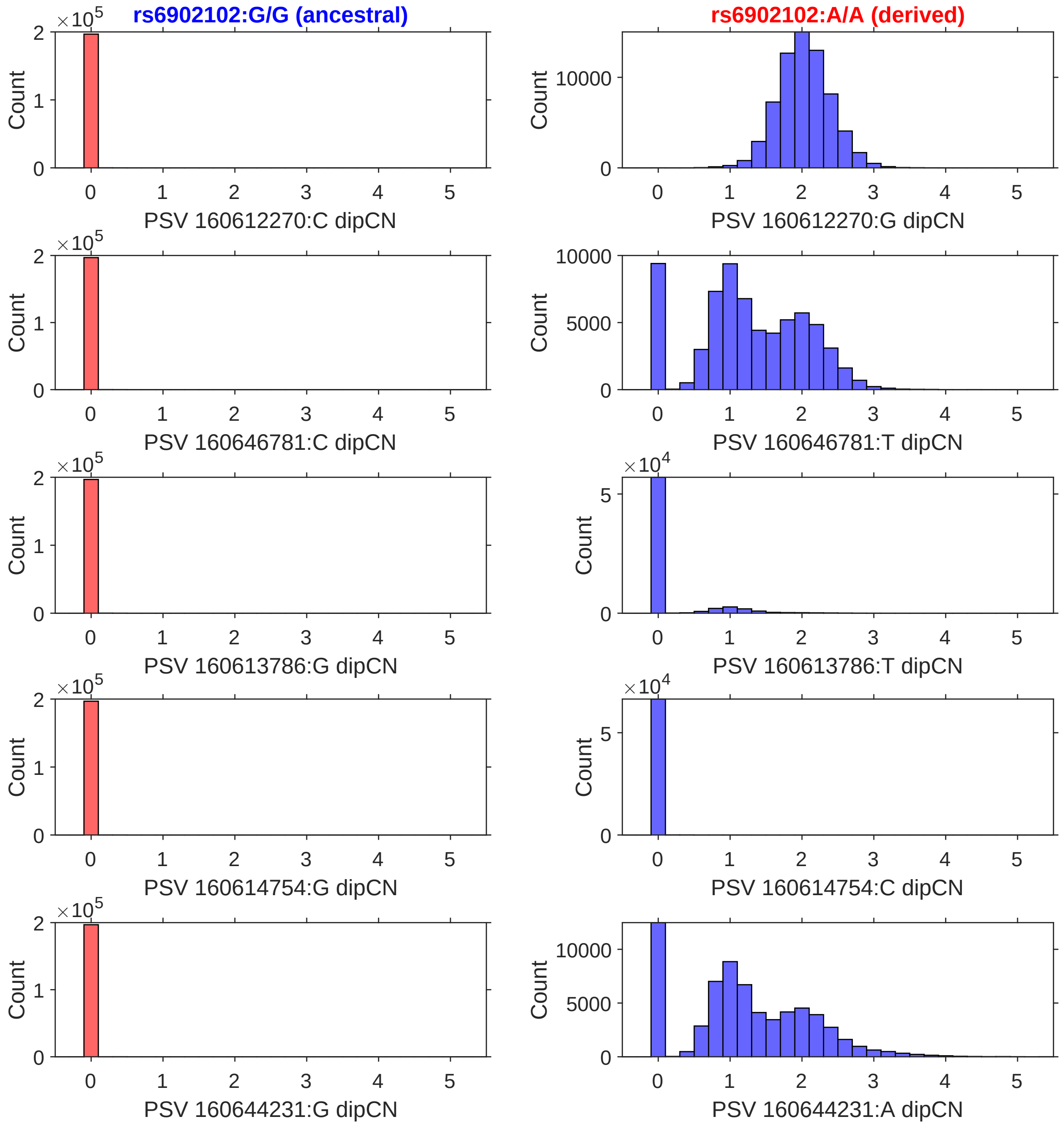


**Supplementary Figure 2. Near-perfect intra-allelic homogeneity of five haplogroup-informative paralogous sequence variants within KIV-2 alleles.** Each histogram shows the distribution of the estimated number of KIV-2 repeat units (summed across an individual's two haplotypes) that contain the "wrong" PSV allele. Left: alleles in UKB participants homozygous for the major KIV-2 haplogroup (based on tag SNPs in tight LD with rs6902102 flanking the KIV-2 repeat; Supplementary Note 3) contain REF alleles of all five PSVs in all repeat units; PSV ALT alleles have copy number 0 (with extremely rare exceptions). Right: alleles in individuals homozygous for the minor KIV-2 haplogroup contain few or no PSV REF alleles. The exceptions follow predictable patterns (Fig. 1D): PSV 160612270:G appears in just the first repeat unit of each allele; PSV 160646781:T is often found in the last repeat unit; PSV 160613786:T is occasionally found in the first repeat unit; PSV 160644231:A is often found in the last 1–2 repeat units.

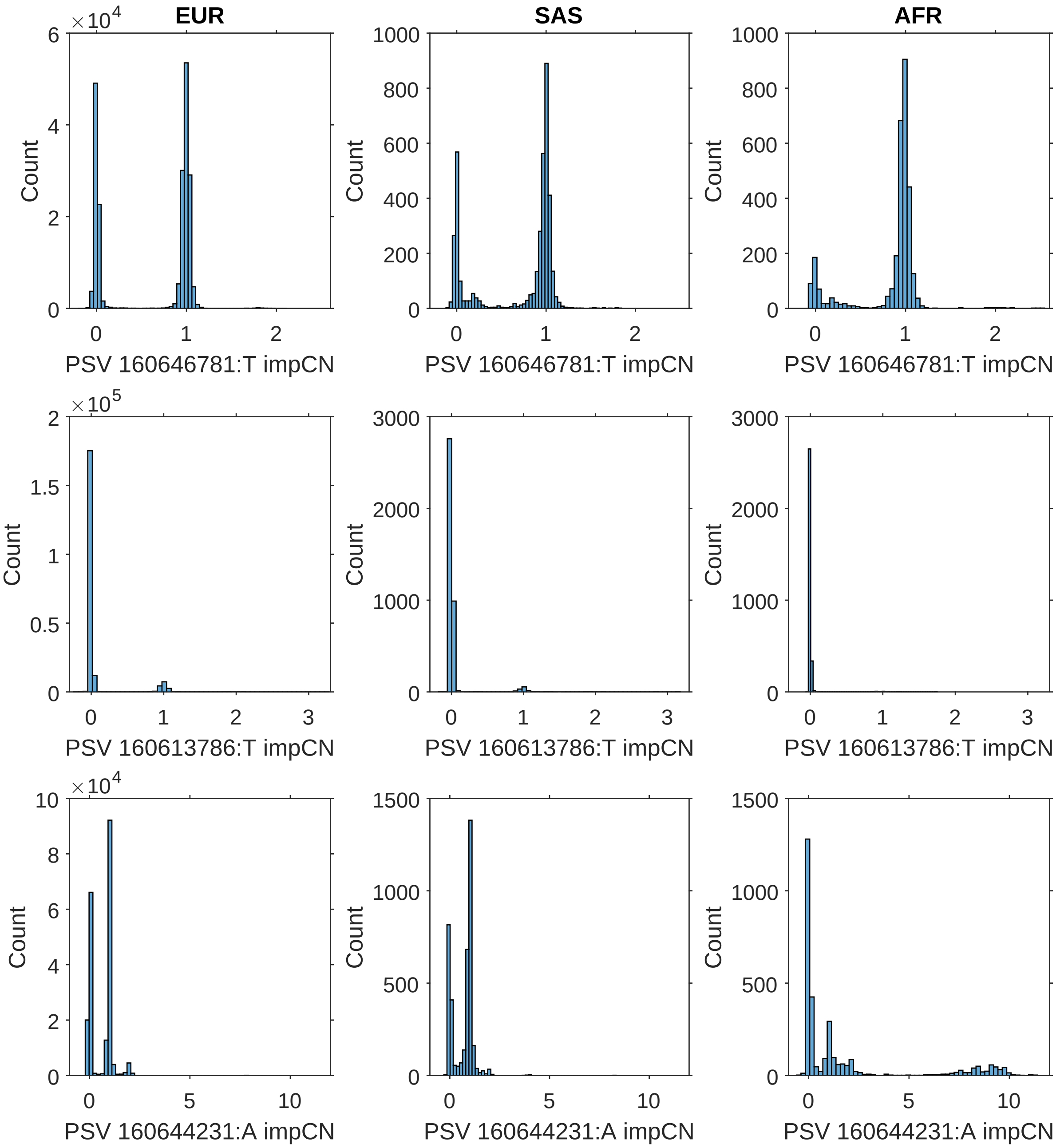


**Supplementary Figure 3. Exceptions to intra-allelic homogeneity of KIV-2 haplogroup-informative PSVs can be accurately imputed from SNP haplotypes.** For the three haplogroup-informative PSVs with common variation in the number of repeat units (of KIV-2 alleles belonging to the derived haplogroup) in which the PSV carries the "wrong" allele, we used SNP haplotypes to impute the number of such exceptions per allele, thereby enabling use of these PSVs when estimating phased KIV-2 allele lengths (Methods). The histograms show the imputed copy numbers of these exceptions in derived-haplogroup alleles of UKB participants for whom we performed PSV-based phasing. Imputation confidence was generally very high, with imputed copy numbers of exceptions typically near integers. For PSV 160644231:A, some alleles contain several exceptions (bottom right), so we did not use this PSV for KIV-2 length estimation if the imputed number of exceptions exceeded 2.5 (as imputation confidence was lower for such alleles).

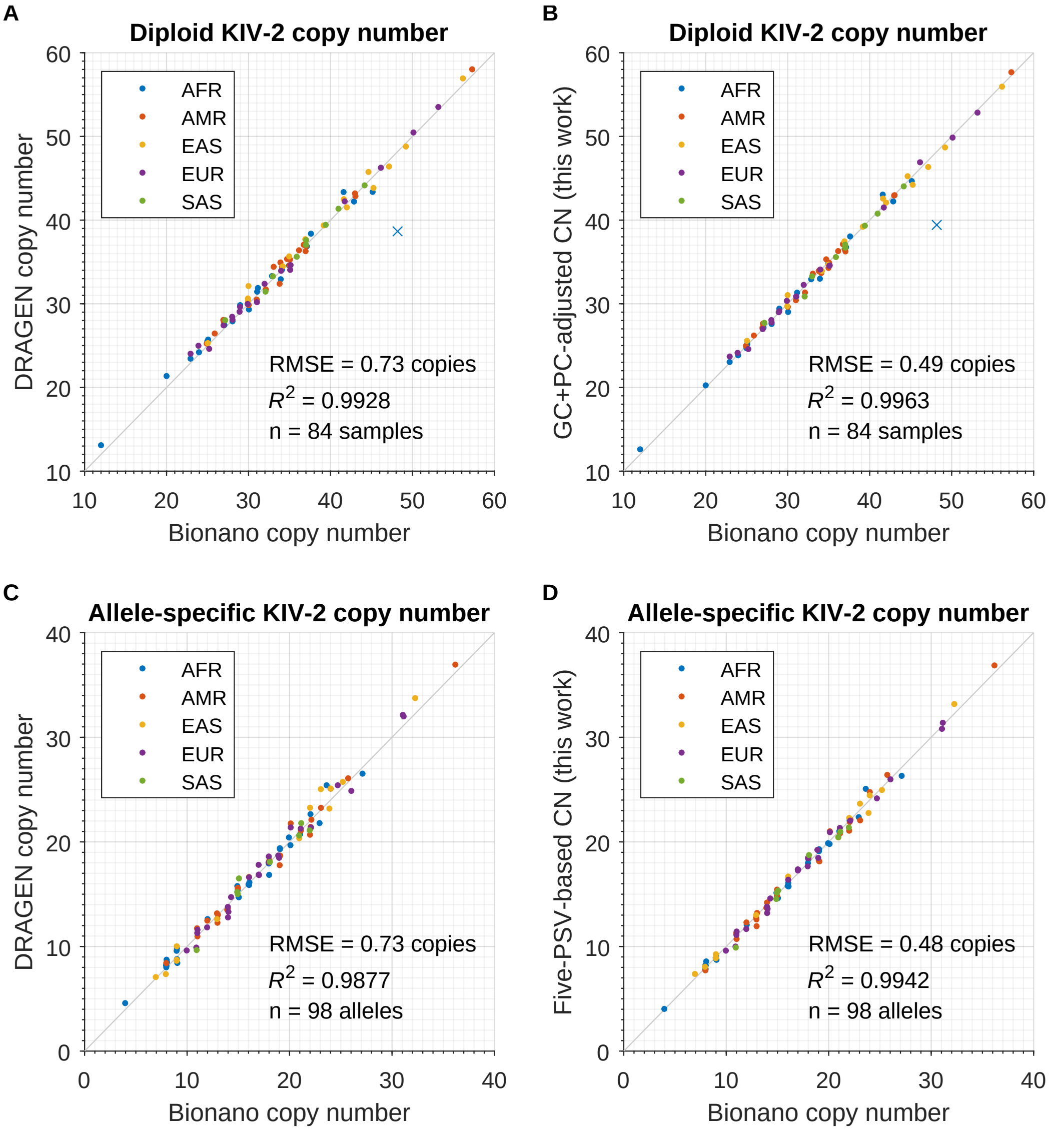


**Supplementary Figure 4. Benchmarks of the DRAGEN KIV-2 CN caller and the method developed here. (A,B)** Benchmarks of diploid KIV-2 copy number estimates on 1000G samples. Data are shown for 85 samples with high-confidence Bionano allele length estimates according to ref.[32]. The blue '×' indicates one outlier (HG02817) probably arising from error in the Bionano estimates, as neither reported Bionano allele length (∼21 and ∼27 copies) is consistent with either allele of HG02818 (a child of HG02817, with allele lengths of 16 and 25 copies in HGSVC3 assemblies[39], consistent with estimates of 15.74 and 24.83 copies from our WGS-based analysis of HG02818). As such, we excluded HG02817 from the benchmark. **(C,D)** Benchmarks of allele-specific KIV-2 copy number estimates on a subset of 49 samples (98 alleles) heterozygous for the two KIV-2 haplogroups (ancestral/derived) with two well-separated alleles (differing by 3+ copies, such that the longer WGS-derived allele length could be assumed to correspond to the longer Bionano allele length). Bionano allele lengths align vertically near integer gridlines, demonstrating their gold-standard accuracy. The RMSE values obtained by our method in 1000G are slightly larger than in UKB (RMSE=0.4 copies), as expected given the smaller 1000G cohort size (which made PCA-based denoising and imputation of PSV exception counts more difficult; Supplementary Note 4).

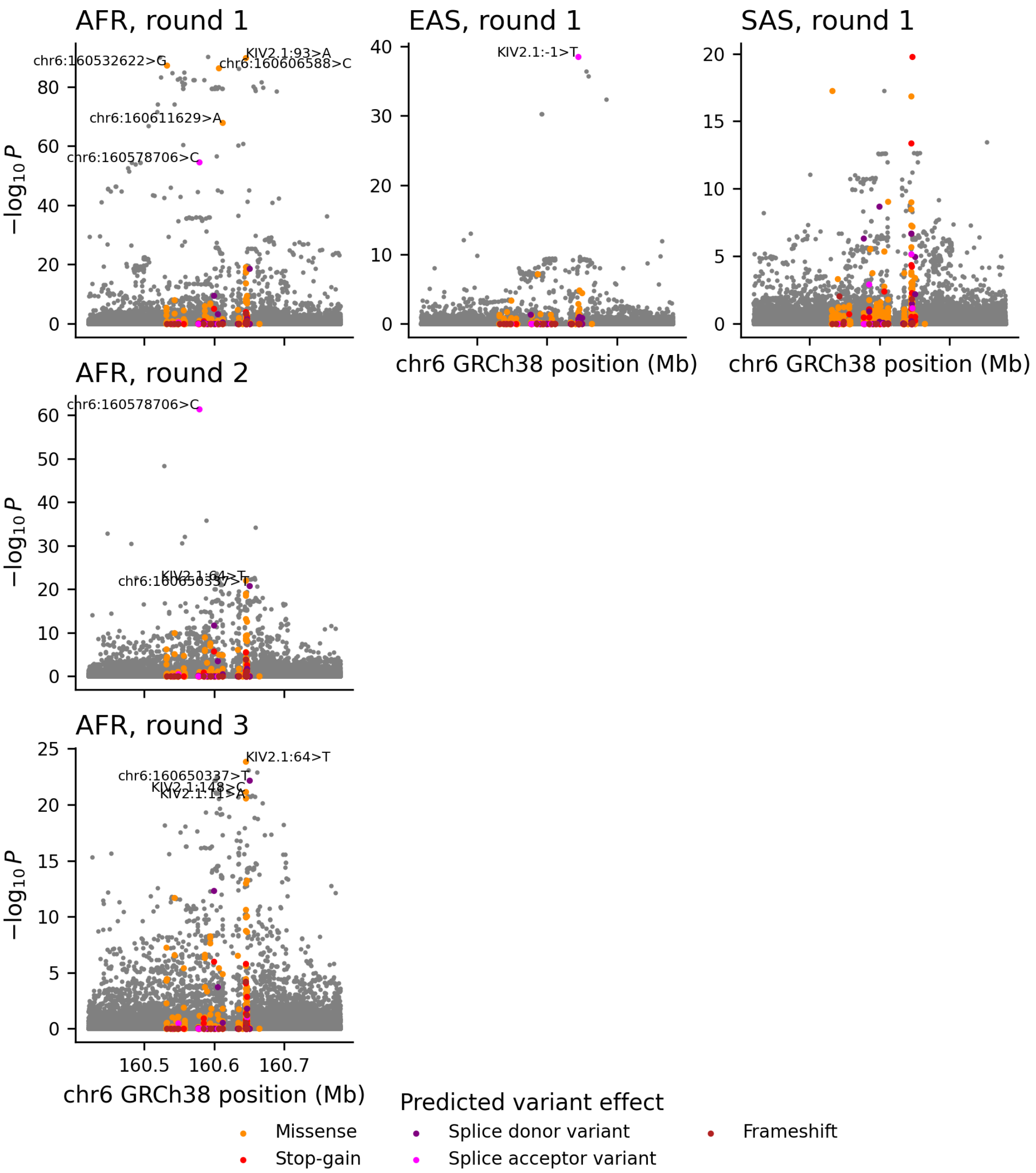


**Supplementary Figure 5. Fine-mapping of key ancestry-specific Lp(a)-modifying variants.** Manhattan plots displaying strength of association ($-\log_{10} P$) versus genomic location for variants near *LPA* are provided for each round of fine-mapping. Columns of plots correspond to fine-mapping in different non-EUR genetic ancestries represented in UKB. Colors indicate variant consequences annotated by VEP. In each round of analysis, we used linear regression to test each variant for association with the difference between measured and predicted $\log_{10}$ Lp(a), cropping low Lp(a) values at 3.8 nmol/L (Methods). In round 1, predictions were based on variants implicated by our previous fine-mapping analysis[26]. In round 2, we included KIV2.1:93>A and KIV2.1:–1>T in the Lp(a) model; in round 3 we included KIV2.1:93>A, KIV2.1:–1>T, and chr6:160578706>C.

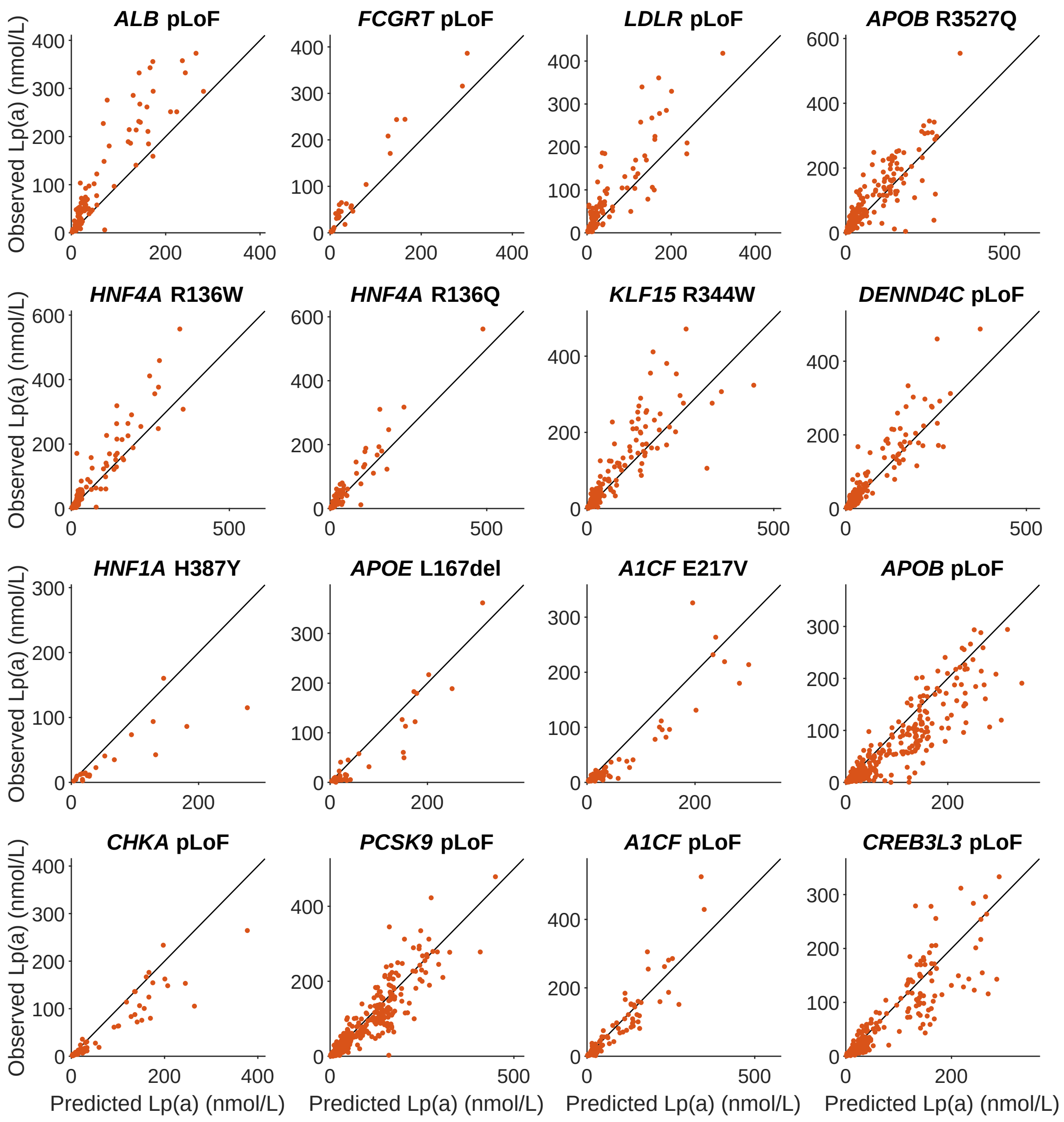


**Supplementary Figure 6. Rare coding variants associated with large effects on Lp(a).** Scatter plots of measured Lp(a) versus Lp(a) predicted from *LPA* diplotypes are shown for UKB EUR-ancestry heterozygous carriers of rare coding variants that associated with *LPA*-adjusted Lp(a). Eight of the plots show data for merged pLoF variants in genes that reached $P<10^{-6}$ significance in pLoF burden tests (eight genes); the other eight plots show data for individual rare coding variants that (i) reached $P<5\times10^{-8}$ significance in single-variant association tests, (ii) had effect sizes of at least 0.5 standard deviations on *LPA*-adjusted Lp(a), and (iii) either affected genes not in the pLoF burden $P<10^{-6}$ set or had effect directions opposite to pLoF effects (eight variants).

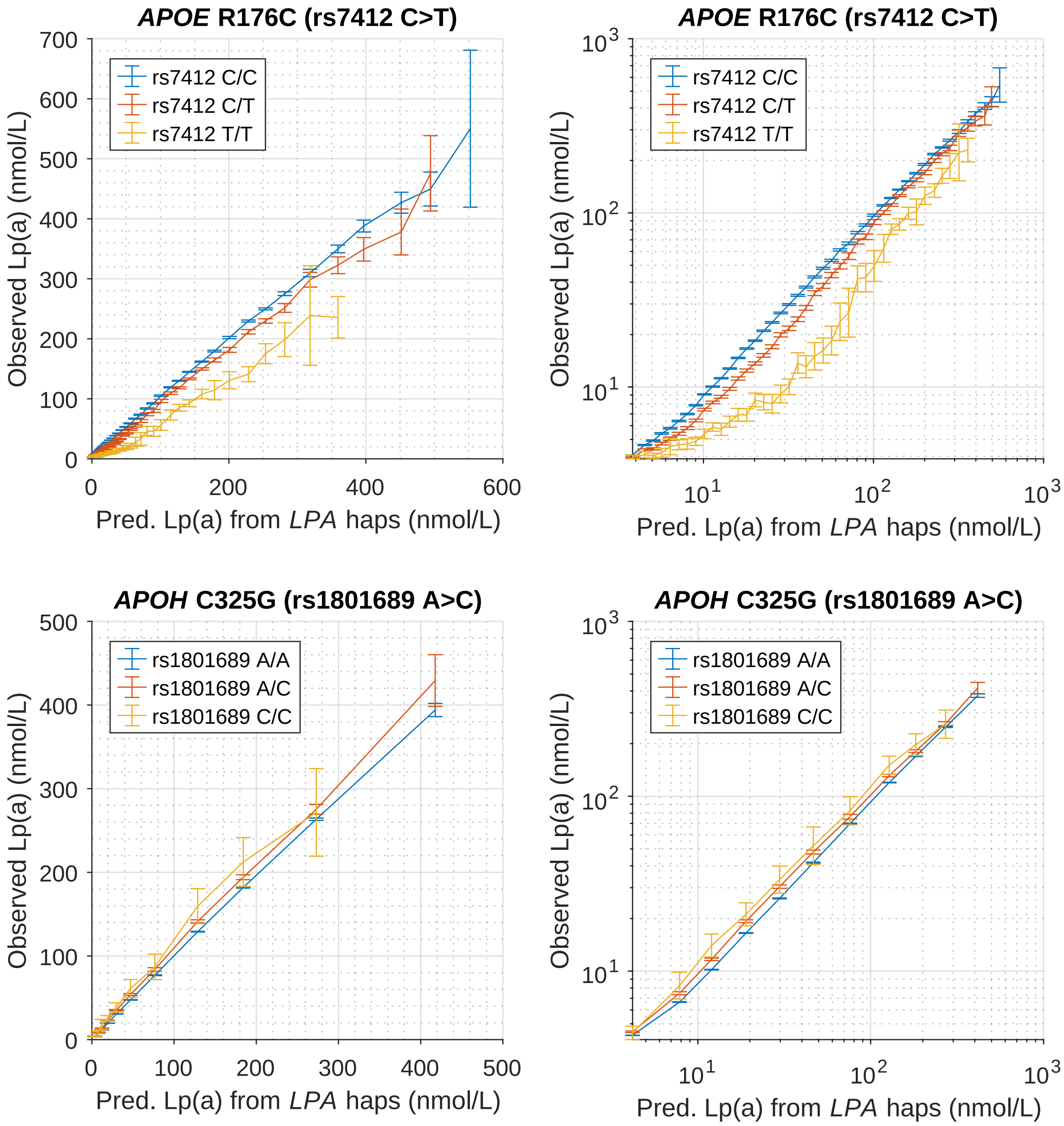


**Supplementary Figure 7. Genetic interactions of *APOE* and *APOH* missense variants with *LPA* haplotypes.** In each plot, observed Lp(a) measurements are compared to genetic predictions of Lp(a) (based on our model of KIV-2 length and sequence variants in *LPA*) in UKB EUR-ancestry participants, stratified by *APOE* rs7412 (*APOE*-$\epsilon$2) genotype (top) or *APOH* rs1801689 (C325G) genotype (bottom). Log-log plots are provided in the right two panels. Both the *APOE* and the *APOH* missense variants appear to have modifier effects on Lp(a) that are approximately multiplicative with the effects of *LPA* haplotypes; however, *APOE*-$\epsilon$2 homozygotes deviate somewhat from this model. Error bars, 95% CIs.

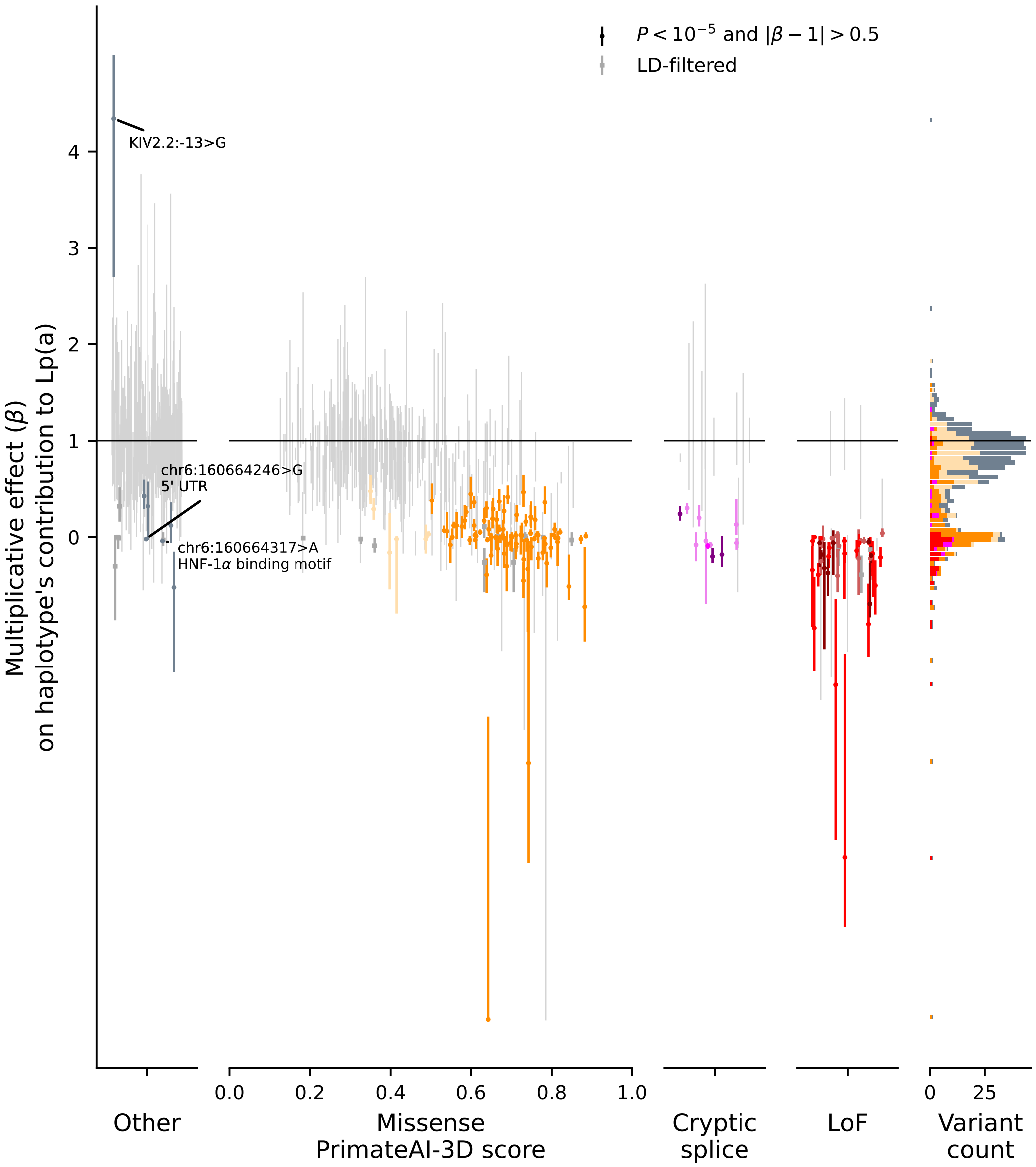


**Supplementary Figure 8. Estimated multiplicative effect sizes of rare variants in *LPA* on Lp(a) produced by an allele.** (left) Maximum likelihood estimate of each rare variant's effect on a haplotype's contribution to Lp(a). (right) Histogram of estimated effect sizes. Color indicates functional annotation as in Fig. 5A. In the left panel, variants with association data that did not satisfy the criteria for likely-causal large effects on Lp(a) are plotted in light grey. Error bars, span of the $\exp(-1.96^2/2)$-likelihood region (approximating a 95% CI). The association of KIV2.2:–13>G with increased Lp(a) (top left) appears unlikely to be causal: it was driven by higher-than-predicted measurements from seven individuals whose alternate alleles segregated with the cryptic splice variant KIV2.2:182>T (G4925A), whereas Lp(a) measurements for the remaining carriers (two EUR and three AFR-ancestry individuals) were consistent with predictions (0.93 mean (measured Lp(a) adjusted for opposite haplotype / predicted Lp(a)), s.e.=0.25).

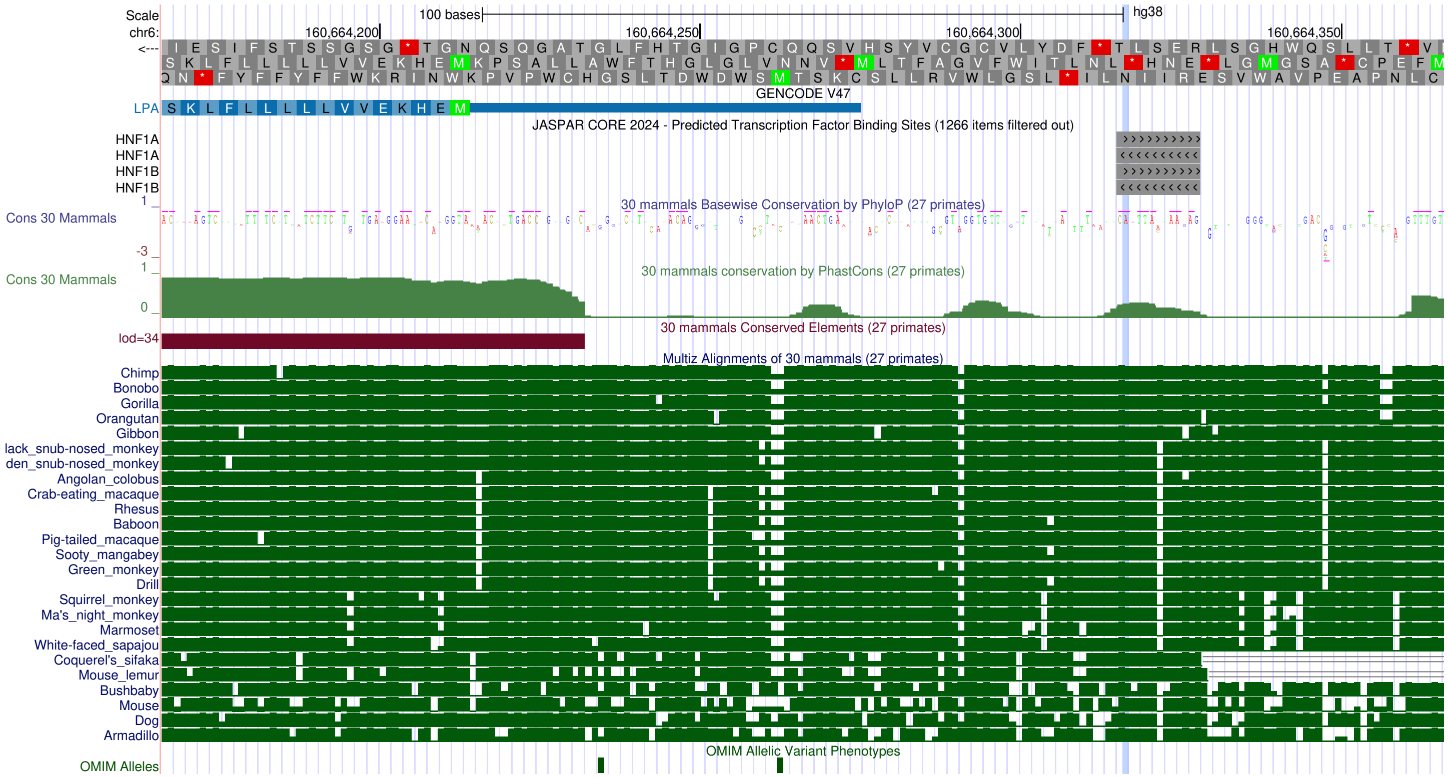


**Supplementary Figure 9. Sequence context and conservation of *LPA* promoter variant rs1320516092 associated with loss of the allele's Lp(a) production.** UCSC Genome Browser[132] screenshot displaying the location of Lp(a)-associated promoter variant chr6:160664317>A (blue highlight), locations of JASPAR[133]-predicted HNF-1$\alpha$ and HNF-1$\beta$ binding sites, and Multiz[134] multiple sequence alignment and conservation scores across 27 primates.

**Supplementary Table 1.** Haplogroup-informative PSVs for KIV-2 repeat phasing.

**Supplementary Table 2.** Lp(a) model parameters for KIV-2 repeat alleles.

**Supplementary Table 3.** Lp(a) model parameters for SNPs, PSVs and indels.

**Supplementary Table 4.** Index variants with statistically independent associations with *LPA*-adjusted Lp(a).

**Supplementary Table 5.** pLoF burden associations with *LPA*-adjusted Lp(a).

**Supplementary Table 6.** Large-effect associations of rare coding variants with *LPA*-adjusted Lp(a).

**Supplementary Table 7.** Lp(a) associations of rare coding, splice region, and promoter variants in *LPA*.
